# TRPV4 Inhibition Reduces Cartilage Growth During Axolotl Limb Regeneration

**DOI:** 10.1101/2025.06.10.658903

**Authors:** Vineel Kondiboyina, Melissa Miller, Maren Ritterbuck, Ashlin Owen, Timothy J Duerr, James R Monaghan, Sandra J Shefelbine

**Affiliations:** Department Of Bioengineering, Northeastern University, Boston, MA; Department of Biology, Northeastern University, Boston, MA; Department Mechanical and Industrial Engineering, Northeastern University, Boston, MA; Institute for Chemical Imaging of Living Systems, Northeastern University, Boston, MA, United States

**Keywords:** Mechanosensitive ion channels, Chondrogenesis, Mechanotransduction, Calcium Signaling, Blastema Growth, Cartilage Patterning

## Abstract

**Background:** Axolotls can regenerate entire limbs, recapitulating mammalian developmental processes. During development, mechanosensitive ion channels such as TRPV4, PIEZO1, and PIEZO2 regulate tissue morphogenesis by transducing mechanical signals. Their roles in regeneration, however, have yet to be thoroughly explored. To investigate this, we assessed the expression of these channels during limb regeneration using single-cell RNA sequencing dataset, hybridization chain reaction fluorescence in-situ hybridization, and immunofluorescence. Additionally, functional relevance was tested by pharmacological inhibition of TRPV4 and PIEZO1/2 mechanosensitive ion channels during limb regeneration.

**Results:** While PIEZO1 expression was undetected, we observed TRPV4 and PIEZO2 expression in uninjured cartilage and at both the mid-bud and palette blastemal stages of limb regeneration. TRPV4 and PIEZO2 were highly expressed in chondrocytes, with PIEZO2 enriched in fibroblasts. Inhibition with GSK205, a TRPV4 antagonist, significantly reduced calcium influx and humeral length without inhibiting cartilage differentiation. Treatment with gadolinium chloride, a broad spectrum mechanosensitive ion channel inhibitor, had no significant morphological impact.

**Conclusions:** TRPV4 and PIEZO2 are dynamically regulated during axolotl limb regeneration. Selective TRPV4 inhibition altered final limb morphology, but chondrogenesis was unaffected. This suggests a role for these genes in shaping tissue architecture during limb regeneration. These findings underscore the importance of ion channel–mediated mechanotransduction in regenerative patterning.

**Key findings:**

- TRPV4 and PIEZO2 show distinct, stage- and cell type-specific expression patterns during axolotl limb regeneration
- Selective pharmacological inhibition of TRPV4 significantly reduces calcium signaling in chondrocytes and leads to shorter limb and humeral lengths in vivo

## 1. Introduction

The role of mechanics in skeletal development is pivotal, with extensive research elucidating the significance of mechanical factors during this process^1–3^. Chondrogenic progenitor cells, which are fundamental for endochondral ossification, perceive their mechanical surroundings through cell– cell and cell–extracellular matrix adhesion, mechanosensitive ion channels, and primary cilia^4^. Among these, mechanosensitive ion channels play a crucial role in transducing mechanical stimuli into biochemical signals, influencing cellular fate and tissue organization.

A key mediator of mechanotransduction is calcium signaling, a ubiquitous secondary messenger that governs diverse biological processes, ranging from embryogenesis to apoptosis^5,6^. Among the many processes calcium signaling is involved in is skeletal differentiation during limb development, where it is essential for endochondral ossification of limb skeletal elements^7^. Mechanosensitive ion channels serve as critical regulators of calcium flux, allowing cells to dynamically adjust their behavior in response to mechanical cues^8–10^. Two major classes of mechanosensitive ion channels involved in skeletal development are the transient receptor potential vanilloid 4 (TRPV4) channel and the PIEZO family of channels. TRPV4, a calcium-permeable non-selective cation channel, is known to mediate responses to osmotic and mechanical stimuli^11,12^, playing a key role in chondrocyte differentiation^13^ and cartilage homeostasis^14^. Moreover, cell volume expansion mediated by TRPV4 activation, has been shown to regulate osteogenesis in stem cells embedded in 3D hydrogels^15^. PIEZO channels, particularly PIEZO1 and PIEZO2, are large, stretch-activated ion channels that facilitate calcium influx in response to mechanical deformation of the cell membrane. These channels are instrumental in mechanosensation and have been shown to influence various developmental processes, including osteoblast differentiation in mesenchymal stem cells^16–18^.

A limitation to the study of mechanosensitive ion channels is that most experiments to date have been conducted in-vitro^13,14,16^. While the in-vitro manipulation of stem cell differentiation by mechanical cues holds promise in tissue engineering and regenerative medicine^4^, the in-vivo role of mechanosensitive ion channels during limb development and regeneration, remains less understood. The few in-vivo studies that do exist have primarily focused on limb development rather than regeneration^17–19^, leaving a significant gap in our understanding of how mechanical forces influence tissue regeneration after injury.

Among vertebrates, the axolotl (*Ambystoma mexicanum*) is a unique model to investigate these questions. Unlike traditional mammalian models such as mice, which are limited to digit tip regeneration, axolotls can fully regenerate entire limbs throughout adulthood^20,21^. Furthermore, the regenerating limb is highly amenable to in-vivo experimental manipulation. Importantly, axolotl limb regeneration recapitulates many molecular and morphogenetic features of embryonic limb development^22–24^, yet it does so in the context of pre-existing tissues, and injury-induced signaling^20,23,25,26^. Limb regeneration in axolotls proceeds through the formation of a blastema, a transient population of dedifferentiated, lineage-restricted progenitor cells that forms at the stump of an amputated limb and orchestrates regrowth after limb amputation^22,24^. These dedifferentiated mesenchymal cells can then activate a development-like transcriptional program to facilitate skeleton reformation. These distinctions make axolotl regeneration an exceptional model for disentangling how mechanical cues, particularly those mediated by mechanosensitive ion channels, govern dedifferentiation and skeletal re-patterning. Despite this potential, how mechanical cues transduced by ion channels influence regeneration is largely unknown. Understanding these differences is essential for improving our understanding of skeletal regeneration and advancing regenerative medicine.

While several studies have elucidated the critical role of calcium dynamics in skeletal development and maintenance^14,17,18^, our understanding of their role in the context of skeletal regeneration remains limited. A recent study showed that inhibiting voltage-gated calcium channels in regenerating axolotl tail inhibits regeneration, emphasizing the significance of calcium ion kinetics in regeneration^27^. Furthermore, perturbing TRPV4 alters joint shape during axolotl limb regeneration^28^. However, this treatment was administered during joint formation, after the blastema cells have differentiated rather than during the earlier blastemal stage before blastema cells commit to a chondrogenic lineage^29^. While these studies provide valuable insights into the impact of mechanotransduction during joint cavitation, the role of TRPV4 and PIEZO1/2 in blastemal cells and chondrogenic progenitors in the regenerating limb has yet to be explored.

This study aimed to bridge this critical knowledge gap by mapping the spatiotemporal expression of TRPV4, PIEZO1, and PIEZO2 across different stages of axolotl limb regeneration at both the transcriptional and protein levels. Furthermore, we investigated the functional role of these mechanosensitive ion channels in vivo by inhibiting their activity during blastemal growth. Our findings reveal that TRPV4 and PIEZO2 exhibit distinct, stage-specific expression patterns and that TRPV4 inhibition perturbs skeletal morphology in the final regenerate without disrupting overall regeneration. These results provide new insights into how mechanosensitive ion channels contribute to tissue patterning and regenerative growth.

## 2. Results

### 2.1. Expression of Mechanosensitive Ion Channels During Axolotl Limb Regeneration

The first objective of this study was to evaluate the spatiotemporal expression of *Trpv4, Piezo1*, and *Piezo2* in regenerating axolotl limbs. Given their known role in shaping skeletal morphology, we hypothesized that these genes would be expressed in fibroblasts and cartilaginous precursors in the regenerating limb.

To achieve this, we reanalyzed existing single-cell RNA sequencing (scRNA-seq) data and performed both Hybridization Chain Reaction Fluorescence In-Situ Hybridization (HCR-FISH) and immunofluorescence to assess *Trpv4*, *Piezo1*, and *Piezo2* expression at the mid-bud and palette stages of limb regeneration. We additionally performed these analyses on uninjured limbs to assess the expression of these genes in articular cartilage to understand how they change following injury. This comprehensive approach captured the dynamic changes in the spatiotemporal expression of mechanosensitive ion channels throughout the limb regeneration process.

#### 2.1.1. Single Cell RNA Sequencing Reveals Distinct, Cell-Type and Stage-Specific Expression Patterns of *Trvp4*, *Piezo1*, and *Piezo2* During Axolotl Limb Regeneration

We first re-examined a previously published dataset^30^ to identify which cell types express *Trpv4*, *Piezo1*, and *Piezo2* in uninjured limbs and in mid-bud and palette staged blastemas.

Cartilage cells in the uninjured limb exhibit the highest *Trpv4* expression, while some expression can be observed in the fibroblast and epidermal populations^8^ (figure 1). At the mid-bud stage, *Trpv4* expression can be found in many populations, including immune, secretory, epidermal, endothelial, cartilaginous, and fibroblast populations. By the palette stage, *Trpv4* expression is notably enriched in the epidermal and cartilaginous populations. Importantly, *Trpv4* expression appears to increase in cartilaginous cells from the mid-bud to the palette stage. *Trpv4* expression was highest in uninjured limbs, potentially indicating that mechanotransduction becomes increasingly more important as chondrocytes mature. While *Trpv4* appeared to be more lowly expressed in the blastema fibroblasts, *Piezo2* expression was enriched in blastema fibroblasts. Fibroblast-specific *Piezo2* expression peaked at the mid-bud stage and was reduced at the palette stage and in uninjured limbs. This expression in fibroblasts appears to be anticorrelated with *Trpv4* expression in cartilaginous cells, suggesting that mechanotransduction during limb regeneration could operate via *Piezo2* in early blastema cells, but then transition to *Trpv4* as fibroblasts begin to differentiate into chondrocytes. Moreover, by the palette stage, *Piezo2* expression is further elevated in cells of the wound epidermis. We observed that *Piezo1* displays broad, non-specific expression across all tissue types and time points, with the highest levels detected in endothelial cells of uninjured limbs. See supplementary material (figure S1) for the cumulative gene expression of the ion channels in limb tissues across all regeneration timepoints.

**Figure 1.**
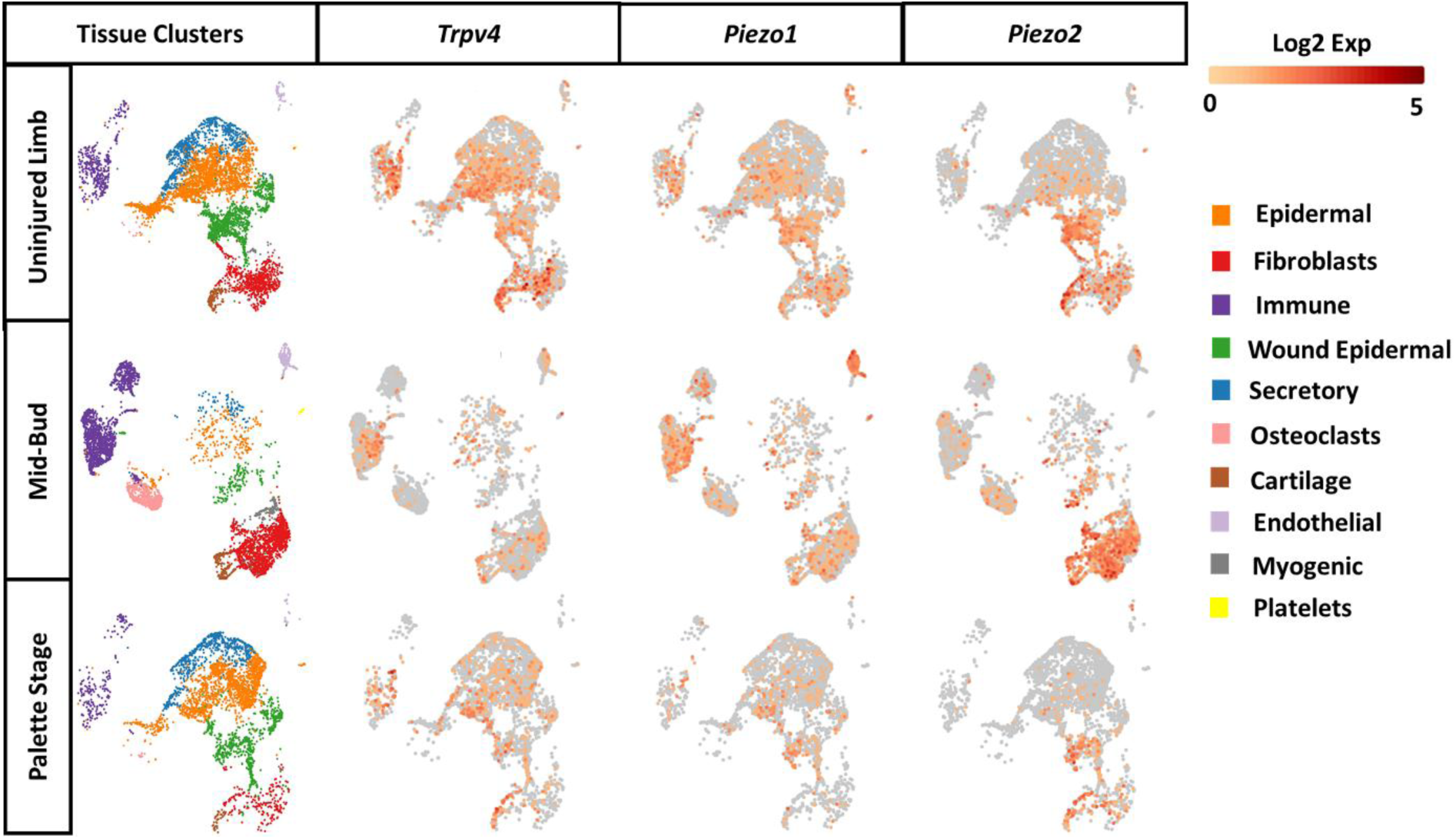
Single Cell RNA data organized by regeneration timepoint, and *Trpv4*, *Piezo1,* and *Piezo2* expression.

#### 2.1.2. HCR-FISH and Immunofluorescence Reveal Spatially Distinct TRPV4 and PIEZO2 Expression in Blastemal and Condensing Mesenchymal Cells During Axolotl Limb Regeneration

After confirming that *Trpv4* and *Piezo1/2* were expressed in the cell types of interest, we performed HCR-FISH to visualize the spatial distribution of *Trpv4* and *Piezo1/2* mRNA in regenerating axolotl limbs. Regenerating limbs were collected at mid-bud and palette blastemal stages, and expression of *Trpv4*, *Piezo1*, and *Piezo2* was visualized and compared to that in articular cartilage of uninjured limbs.

We observed high levels of *Trpv4* and *Piezo2* expression in the articular cartilage of uninjured limbs (see figure 2 a-h, negative controls: figure S2-S3). While *Piezo1* expression was not observed in blastemal tissues or uninjured articular cartilage (see figure S4), *Piezo1* expression was detected in the tissue surrounding the humeral joint at the uninjured timepoint (see figure S4 a-h). The absence of robust *Piezo1* expression in uninjured cartilage, and its low expression in the surrounding connective tissue, aligns with scRNA-seq data, confirming its limited presence in blastema and cartilage tissues.

**Figure 2.**
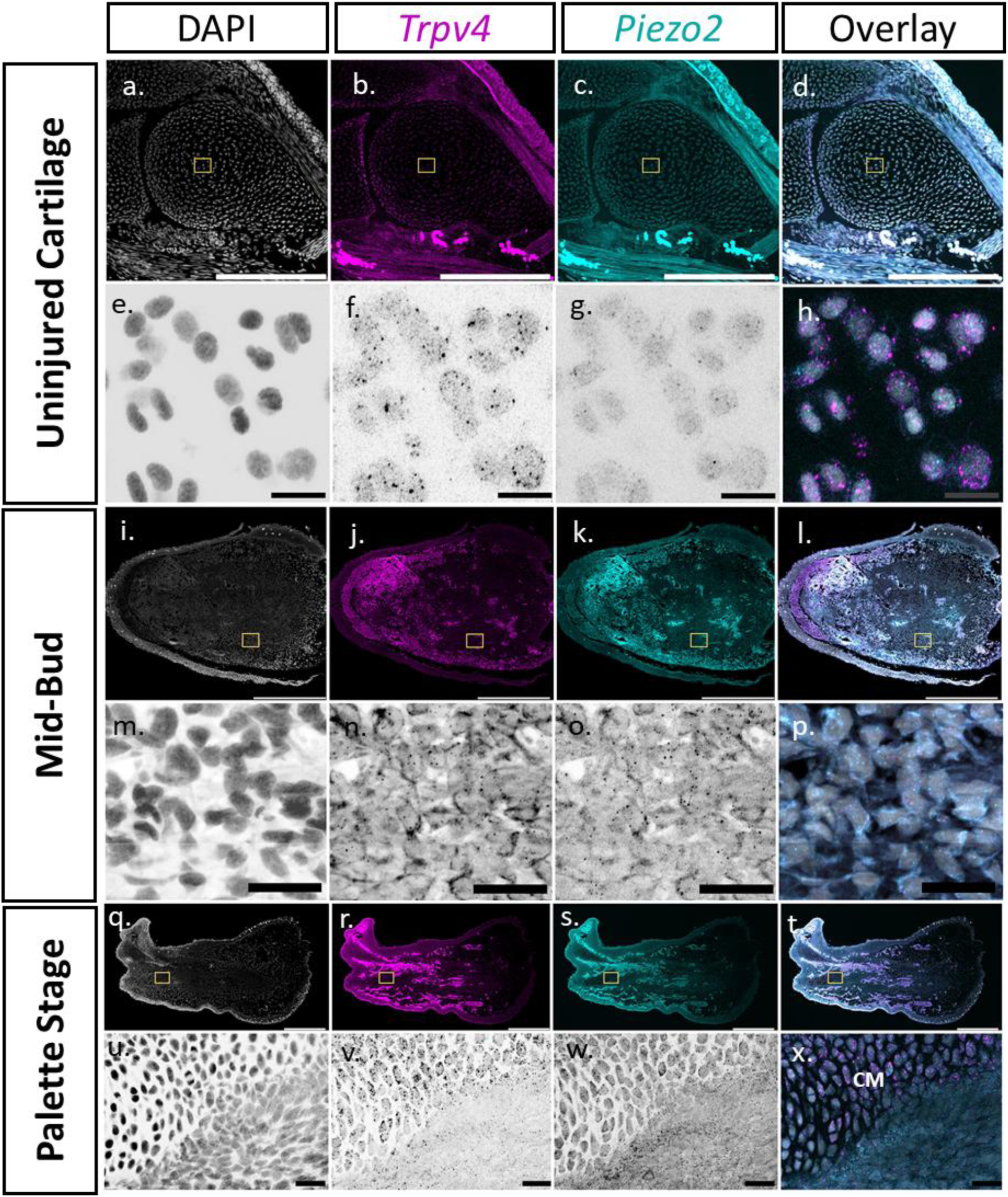
mRNA expression of *Trpv4* and P*iezo2* during limb regeneration (n=3/timepoint) at (a-h) Uninjured cartilage. Scale: (a-d) 500µm (e-h): 25µm (i-p) Mid-bud blastema stage. Scale: (i-l) 1000µm, (m-p) 50µm (q-x) Palette stage blastema. Scale: (q-t) 1000µm (u-x) 50µm. CM: Condensing Mesenchyme and Yellow box represents the region of interest (ROI) and the individual channels in the ROI are visualized as inverted grayscale lookup table images with the darker dots representing respective mRNA while the lighter circular boundary representing the cell nuclei.

At the mid-bud stage, *Trpv4* and *Piezo2* were expressed across the blastemal tissue, with some expression in the epithelium for *Trpv4* (see figure 2 i-p, figure S5). Similarly, at the palette stage, both *Trpv4* and *Piezo2* were expressed in blastemal cells, with the highest expression observed in the condensing mesenchyme or differentiating skeletal elements (see figure 2 q-x).

To complement the transcriptional and spatial data, we employed immunofluorescence to assess protein-level expression and localization of TRPV4 and PIEZO2 at the same timepoints. Given the negligible *Piezo1* mRNA levels, PIEZO1 protein was not analyzed. Consistent with the mRNA data, TRPV4 and PIEZO2 proteins were detected in articular chondrocytes at the uninjured timepoint (figure 3a–b), with no signal observed in negative controls (figure S6). In regenerating limbs, both proteins were expressed throughout the blastema at mid-bud and palette stages, with notably robust expression in the condensing mesenchyme at the palette stage (figure 3c–f).

**Figure 3.**
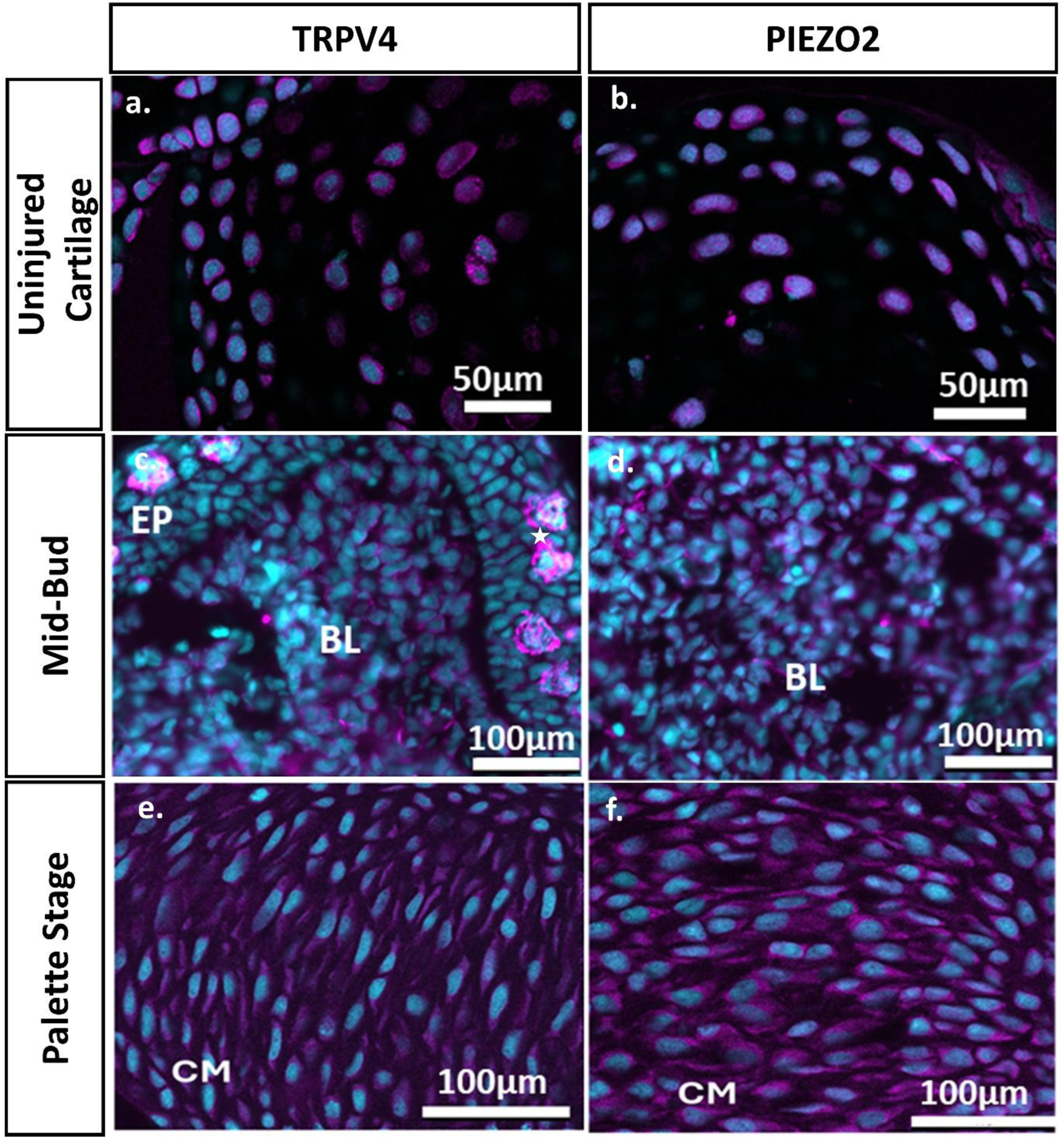
TRPV4 and PIEZO2 immunofluorescence expression during limb regeneration at (a-b) Uninjured cartilage (c-d) Mid-bud stage (e-f) Palette stage blastema. Cyan stains for the nucleus and magenta stains for the presence of TRPV4/ PIEZO2 ion channels. EP: Epithelium, BL : De-differentiated Blastemal cells, CM: Condensed Mesenchyme. White star labels Leydig cells.

Together, these findings confirm the spatial and temporal expression of TRPV4 and PIEZO2 at both mRNA and protein levels during limb regeneration, particularly in regions undergoing mesenchymal condensation, supporting a potential role for these mechanosensitive ion channels in mechanotransduction and skeletal tissue remodeling.

### 2.2. Effect of Mechanosensitive Ion Channel Antagonists on Axolotl Limb Regeneration

After confirming the presence of TRPV4 and PIEZO2 ion channels at both the transcript and protein levels in fibroblasts and cartilaginous cells, we investigated their functional role in axolotl limb regeneration by pharmacologically inhibiting their activity. Specifically, we investigated the impact of GSK205, a selective TRPV4 antagonist, and Gadolinium Chloride (GdCl₃), a non-specific inhibitor of several mechanosensitive ion channels, including TRPV4 and PIEZO1/2^31,32^.

We hypothesized that mechanosensitive ion channels regulate key regenerative processes, such as cellular differentiation and proliferation, and that their inhibition would impair limb regeneration. Accordingly, we expected diminished regenerative capacity in the GSK205- and GdCl₃-treated groups compared to vehicle controls.

Given that mechanosensitive ion channels modulate intracellular calcium dynamics, we first validated the efficacy of the chosen antagonists in-vitro by assessing calcium signaling in axolotl chondrocytes. This ensured that the selected inhibitors effectively disrupted ion channel activity at the intended working concentration. We then applied the same concentration in-vivo to assess the impact of ion channel inhibition on limb regeneration.

#### 2.2.1. Pharmacological Inhibition Of Mechanosensitive Ion Channels Disrupts The Intracellular Calcium Ion Dynamics In Axolotl Chondrocytes In-Vitro

The effects of mechanosensitive ion channel antagonists GSK205 and GdCl_3_, on axolotl chondrocytes were assessed by measuring calcium signaling dynamics under baseline conditions and hypotonic stress. Hypotonic stress is known to increase the percentage of activated chondrocytes compared to baseline conditions and this response is expected to be severely attenuated in the presence of mechanosensitive ion channel antagonists^10,12,33^.

Calcium dynamics of mature in-situ chondrocytes in the distal epiphysis of isolated axolotl humeri were assessed using confocal time-lapse imaging in-vitro. Calbryte® 520-AM calcium indicator-stained samples were pre-incubated in either a DMSO vehicle control, 10 µM GSK205, or 10 µM GdCl_3_ before imaging and remained in the respective solution throughout the experiment. Following an initial recording in baseline conditions, the medium was replaced with a hypotonic solution containing the same vehicle or drug concentration, and imaged to capture the chondrocyte response. Calcium signaling activity across treatment groups, at baseline and after hypotonic activation, was analyzed using previously established protocols^11^. See figure 4 a-b for representative confocal fluorescence images and corresponding calcium signaling trace. Similar to previously published data on mammalian chondrocytes, we observed multiple calcium signaling peaks within each cell during our imaging window of 10 minutes^11^.

**Figure 4.**
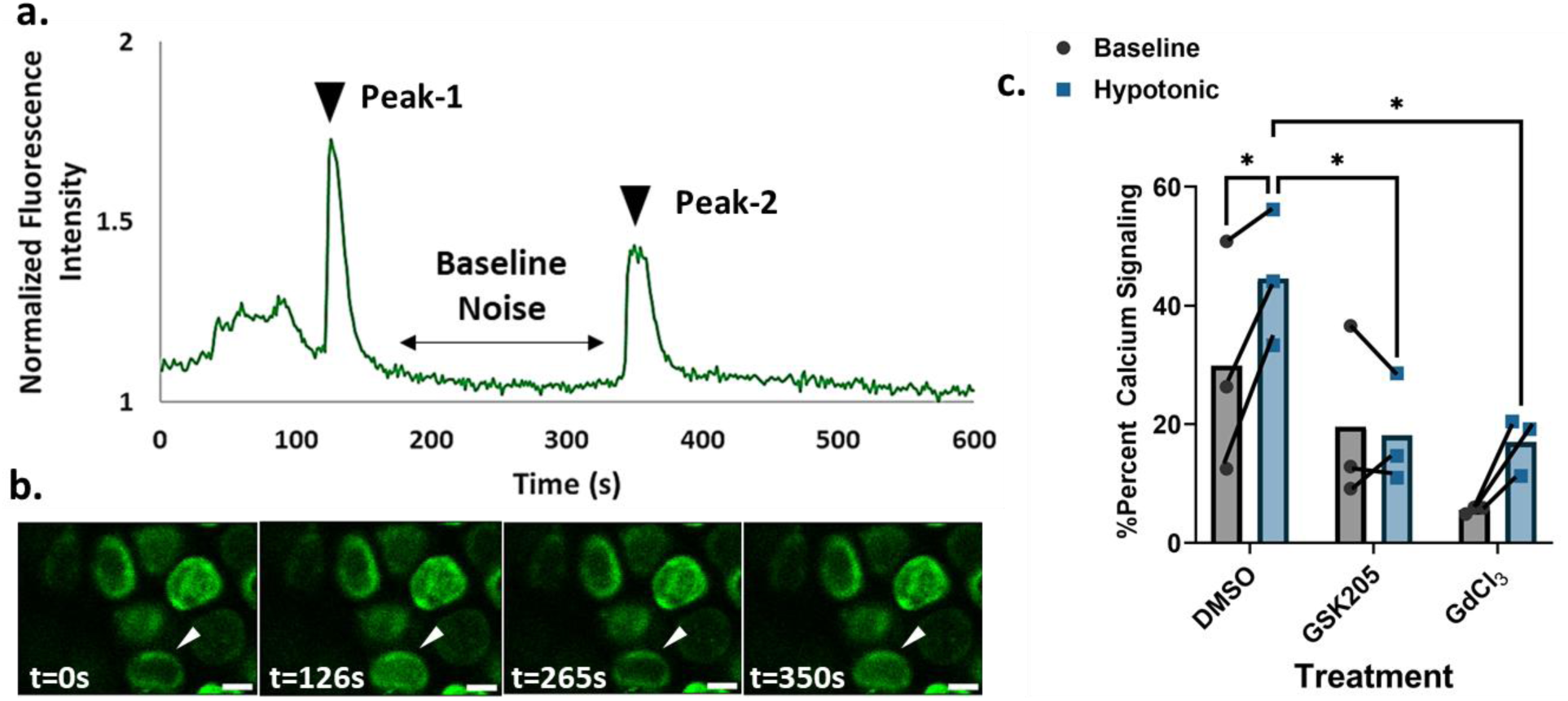
(a) Representative calcium signaling trace of a single chondrocyte in DMSO control group over time. A peak represents a >4x increase in calcium fluorescence intensity compared to baseline fluctuations (b) Representative images of calcium indicator-stained chondrocytes over time with white arrows indicating signaling chondrocytes. Scale 10µm. (c) Percentage calcium signaling in mature chondrocytes before and after hypotonic activation (n=3 per treatment group). * represents statistical significance with p values less than 0.05.

The percentage of signaling cells upon hypotonic activation was significantly elevated compared to the baseline in the DMSO control group (mean change: 14.65% ,p<0.05). Baseline signaling levels were lowered in both GSK205 and GdCl_3_ treated samples, indicating that steady state calcium signaling levels were effectively reduced upon treatment. However, hypotonic activation in GSK205 treated samples did not increase the percentage of signaling cells indicating effective blockage of channel activation (mean change: −1.45%, p=0.8). In contrast, while we observed an increase in percentage calcium signaling upon hypotonic activation in the GdCl_3_ group, it was not statistically significant (mean change: 11.38%, p=0.09). Furthermore, the calcium signaling response was muted in the GSK205 and GdCl_3_ group compared to the control group with the overall percentage of signaling upon activation in GSK205 and GdCl_3_ groups significantly lower than in control group (p<0.05, see figure 4c).

These results taken together show that TRPV4 specific antagonist GSK205 effectively blocked the ion channel activity while GdCl_3_, a non-specific mechanosensitive ion channel antagonist, reduced ion channel response but did not completely abolish it.

#### 2.2.2. Selective TRPV4 antagonism impacts joint morphology without impairing limb regeneration in-vivo

We next assessed the impact of perturbing ion channels during limb regeneration. Forelimbs were amputated along the mid-humerus, and at 9 Days Post Amputation (DPA) animals were housed in water supplemented with GSK205, GdCl_3_, or DMSO-controlled vehicle. This time point was chosen in order to inhibit mechanosensitive ion channels in blastema cells prior to differentiation into cartilaginous precursors. Regenerating blastemas were imaged at 9 DPA, 20 DPA and 27 DPA before being collected and stained with alcian blue to visualize final regenerate morphology. Additionally, total limb length and humeral length were measured across treatment groups.

All animals regenerated their limbs with observable cartilage distal to the amputation site (figure 5 a-b). However, GSK205 treated samples had significantly shorter limbs (mean difference: −31.36%, p< 0.001) with short humerii (mean difference: −35.11%, p< 0.001) compared to the control group. Total limb length (mean difference: 4.70%, p=0.38) and humeral length (mean difference: 8.51%, p=0.18) in GdCl_3_ treated animals appeared identical to DMSO treated animals (figure 5 c-d). These results suggest that selective inhibition of mechanosensitive ion channels can influence skeletal morphology without inhibiting the overall ability to regenerate the limb. Despite its broad-spectrum inhibition of mechanosensitive ion channels, GdCl₃ did not significantly reduce humeral or limb length. Given that GdCl_3_ simultaneously inhibits multiple mechanosensitive ion channels, it is likely that residual TRPV4 activity is present in this group, as seen in calcium signaling results in figure 4.

**Figure 5.**
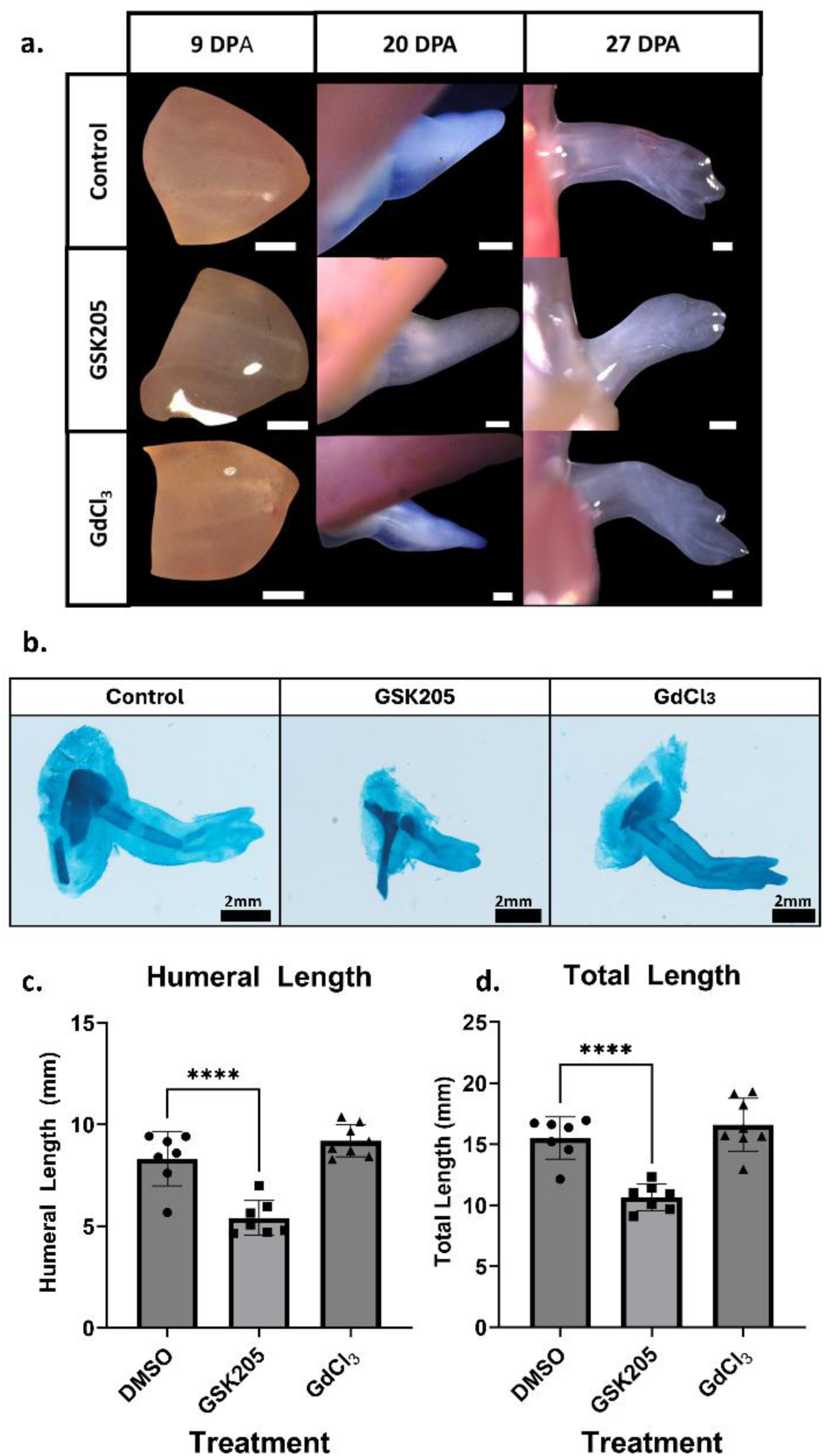
(a). Brightfield images of regenerating blastema undergoing GSK205 and GdCl_3_ drug treatments (n=8 per treatment group). Scale bar: 400µm b. Alcian blue staining of regenerating limbs at the end of the treatment period c. Comparison of humeral lengths across treatment groups at the end of the treatment period. d. Comparison of total limb lengths across treatment groups at the end of the treatment period. **** represent statistical significance with p values less than 0.001.

## 3. Discussion

Mechanosensitive ion channels play fundamental roles in transducing physical forces into biochemical signals, yet their precise contributions to skeletal development and regeneration remain incompletely understood. In this study, we elucidated the spatial and temporal expression patterns of PIEZO1, PIEZO2, and TRPV4 during axolotl limb regeneration and examined their functional relevance in tissue patterning and skeletal development. Our findings revealed that these ion channels exhibited distinct expression patterns during limb regeneration, with TRPV4 emerging as a potential regulator of skeletal growth.

### 3.1. Distinct Expression Profiles of PIEZO1, PIEZO2, and TRPV4 in the Regenerating Limb

Our spatial and single-cell transcriptomic analyses demonstrate dynamic and cell-type-specific expression of mechanosensitive ion channels at key regenerative stages. scRNA-seq analysis at the mid-bud blastema stage showed that *Piezo2* is highly expressed in fibroblasts, which constitute the majority of the blastemal cell population^34^, whereas *Trpv4* is predominantly expressed in the epidermal cells. As regeneration progressed to the mesenchymal condensation stage, HCR-FISH and immunofluorescence analysis demonstrated the co-expression of TRPV4 and PIEZO2 in condensing mesenchyme, implicating these channels in cartilage differentiation and proliferation. This observation aligns with previous studies on limb development, where TRPV4 and PIEZO2 were expressed in condensing mesenchyme during later-stage limb formation in mice^17,35^.

Interestingly, despite the interdigital expression of *Piezo1* in mammalian limb development, a similar pattern was not observed in axolotls during limb regeneration^17^. This disparity could be attributed to variations in interdigital dynamics between mammals and amphibians. Axolotls lack coordinated cell death in the interdigital area, which in mammals is a crucial process for the formation of distinct digits^36^. This mechanistic difference emphasizes the divergence in limb patterning between the two species.

The expression of *Trpv4* in blastemal macrophages as seen in scRNA-seq data raises intriguing questions about their role in immune-regenerative crosstalk. Prior studies indicate that macrophages are essential for axolotl limb regeneration^37^. While our study did not directly assess macrophage depletion, the presence of mechanosensitive ion channels in immune cells suggests that mechanical cues may influence immune-mediated regeneration. Future studies targeting macrophage-specific ion channel activity could provide deeper insights into this interplay.

### 3.2. TRPV4 as a Potential Regulator of Cartilage Growth

While previous studies on limb development highlight TRPV4 as a possible regulator of chondrogenesis^14^, and PIEZO1/2 as modulators of osteogenesis^17^, it is interesting to note that in-vivo studies utilizing *Trpv4* and *Piezo1/2* knockout mice report typical chondrocyte and osteocyte differentiation during limb development albeit with differences in joint morphology, susceptibility to disease or decrease in bone quality^17,19^.

Our previous study on TRPV4 desensitization through TRPV4 agonism during later stages of axolotl limb regeneration showed changes in humeral joint morphology and reduced cellular proliferation^28^. However, since the treatment was initiated post-chondrocyte differentiation, after the joint had already formed, we could not assess the effect of TRPV4 dysfunction on cartilage formation. In this study, we assessed the effects of TRPV4 and PIEZO1/2 inhibition on limb regeneration by treating the animals with mechanosensitive ion channel antagonists GSK205 and GdCl_3_ at the early blastema stage through chondrocyte differentiation and joint formation.

Calcium signaling assays in mature axolotl articular chondrocytes revealed robust mechanosensitive responses to hypotonic stimulation, which were diminished by GSK205 and GdCl_3_ treatment, corroborating prior findings in mammalian chondrocytes^12,19,38,39^. Notably, the GSK205-treated group exhibited no detectable increase in calcium signaling following hypotonic activation, indicating that GSK205 effectively blocked TRPV4 channel activity. This is consistent with previously published data^11^. In contrast, the GdCl₃-treated group displayed a modest but statistically insignificant increase compared to the control group. These findings suggest that while GdCl₃ did not entirely abolish the mechanotransduction of ion channels, it substantially impaired their responsiveness. One of the reasons for reduced potency of GdCl_3_, apart from the non-selective targeting of ion channels, could be that it inhibits channel activation indirectly by binding to membrane phospholipids and increasing local membrane rigidity as opposed to directly interacting with the ion channels^40^.

To assess the impact of ion channel inhibition in-vivo, we imaged regenerating limbs at multiple timepoints and stained the regenerates with alcian blue to examine skeletal morphology. Despite antagonist treatment, regeneration proceeded uninhibited in all groups, with observable cartilage growth and joint formation. However, TRPV4 inhibition using GSK205 led to a significant reduction in both humeral and total limb length, suggesting a role for TRPV4 in cartilage growth. In contrast, combined inhibition of TRPV4, PIEZO1, and PIEZO2 using GdCl₃ did not affect humeral or limb length, despite reduced calcium signaling in this group.

These findings suggest that mechanosensitive ion channels, although expressed in blastema fibroblasts, may not be required for the initiation of regeneration or the early differentiation of blastema cells. We previously identified a proximal-distal stiffness gradient in the late limb blastema associated with skeletal differentiation^41^. Similarly, Edwards-Jorquera et al. showed that differences in matrix stiffness in the proximal vs distal blastema in-vivo with decreased cellular proliferation with increased matrix stiffness in-vitro^42^. Given the strong expression of PIEZO2 in this region, we anticipated a synergistic interaction between matrix stiffness and PIEZO1/2 in guiding fibroblast migration and proliferation. However, our results suggest an alternative mechanism, wherein TRPV4 independently governs cartilage growth and development.

Still, we cannot exclude a role for PIEZO channels during earlier regenerative stages—such as blastema formation or mesenchymal condensation—particularly given the non-specific and potentially suboptimal inhibition provided by GdCl₃. The absence of a phenotype may reflect compensatory mechanisms that preserve skeletal growth under partial inhibition, suggesting that complete or more selective blockade may be necessary to uncover functional roles. Future investigations utilizing genetic or alternative pharmacological approaches^43^ are warranted to dissect the contributions of PIEZO ion channels in axolotl limb regeneration.

### 3.3. Conclusions and Future Directions

Our findings provide novel insights into the mechanosensitive regulation of limb regeneration, revealing distinct expression patterns of TRPV4, PIEZO1, and PIEZO2 across different regenerative stages. While dysfunction of mechanosensitive ion channels does not appear to impede blastemal growth, TRPV4 emerges as a key regulator of cartilage patterning. The incomplete, non-selective inhibition of TRPV4 and PIEZO1/2 did not significantly alter skeletal morphology, suggesting the existence of functionally distinct, threshold-dependent mechanotransduction pathways. These results suggest fundamentally different mechanisms governing blastemal growth and skeletal patterning, highlighting the need for further mechanistic studies to elucidate how mechanosensitive signaling orchestrates limb regeneration. Future research employing genetic tools and selective pharmacological approaches will be essential for uncovering the precise mechanotransduction mechanisms underlying vertebrate appendage regeneration.

## 4. Experimental Methods

All materials and reagents were bought from Thermo Fisher Scientific, Waltham, MA, unless specifically noted otherwise. All animals were bred at Northeastern University, and all procedures and surgeries were approved by the Northeastern University Institutional Animal Care and Use Committee. Surgeries were performed while axolotls were anesthetized in 0.01% benzocaine.

### 4.1. Single Cell RNA Sequencing

Our current investigation involves a reanalysis of scRNA-seq data from Li et al. 2021^30^ with a specific focus on extracting information pertinent to expression patterns of mechanosensitive ion channels during the process of regeneration. The scRNA-seq data was acquired from the upper forearm tissue of 5 axolotls across seven regenerative timepoints, spanning from 3 hours to 33 days post-amputation (DPA), including uninjured control samples. The samples sequenced by Li et al contained the entire blastema in the mid-bud and palette stages and hence proliferating chondrocytes can be identified at these timepoints^30^. The obtained reads were aligned to the axolotl v6 genome, counted, and normalized by read depth using 10X Genomics Cellranger software^45^. The resulting count matrices were then imported into Seurat, where they underwent filtration to retain features expressed in a minimum of five cells. Additional filtering criteria were applied, including a mitochondrial gene content of less than 10%, mRNA counts ranging between 1000 and 20,000, a minimum of 500 expressed features, and a red blood cell gene content of less than 5%.

Normalization of counts was achieved through the application of the Seurat SCTransform() function with regression of mitochondrial gene content^46^. Subsequent to dimensional reduction, the dataset was integrated using the IntegrateLayers() function with Canonical Correlation Analysis (CCAIntegration). A secondary round of dimensional reduction was executed on the integrated data, followed by cluster assignment with a resolution set at 0.6 yielding 10 broad cell type clusters(see Table 1). Expression data and Uniform Manifold Approximation and Projection (UMAP) coordinates were extracted from the integrated Seurat object for visualization with ggplot2^47^.

**Table 1.** Genetic markers used for identification of the cell types.

| Cluster | Marker | Cluster | Marker |
| --- | --- | --- | --- |
| Epidermal | KRT5<sup>22,59</sup>, KRT12<sup>30,60</sup>,<br>KRT17<sup>59,60</sup> | Osteoclasts | CTSK<sup>61</sup> |
| Fibroblasts | PRRX1<sup>22,30</sup>, LUM<sup>22,59,60</sup>,<br>DPT<sup>22,60</sup>, COL1A2<sup>22,30</sup> | Cartilage | COL9A2<sup>24</sup> |
| Immune | MARCO<sup>22,59,60</sup>,<br>IFITM10<sup>62</sup>, ARG1<sup>22,30</sup> | Endothelial | VWF<sup>22,59,60</sup> |
| Basal/Wound<br>epidermal | COL17A1<sup>30,60</sup> | Myogenic | MYF5<sup>30,60</sup>, PAX7<sup>24,60,63</sup> |
| Secretory | FCGBP<sup>59,60</sup> | Thrombocytes | GP1BA<sup>59</sup>, GP1BB<sup>59</sup>,<br>GP9<sup>59</sup> |

### 4.2. Hybridization Chain Reaction Fluorescence In-Situ Hybridization (HCR-FISH)

#### 4.2.1. Animal Handling and Sample Collection

Axolotls (tail to snout length 15-20 cm) were amputated along the mid-humerus. The regenerating limbs from the animals were collected at two distinct time points; 27 DPA representing the mid-bud blastema and 41 DPA representing the palette stage^48^. The uninjured timepoint was harvested from animals with tail to snout length of 8-10 cm. Each timepoint involved the collection of 3 limbs, with each limb sourced from a different animal. Articular cartilage from the uninjured limbs served as a positive control for *Trpv4* and *Piezo1/2* expression^49^. The regenerating blastemas were staged according to Tank et al. (1976) ^48^.

Despite the bigger size of the animal in this study compared to those used in scRNA-seq, it is anticipated that the cell identities at the mid-bud, palette, and uninjured time points remain consistent. The divergence lies in the regeneration time, with smaller animals (8-10 cm) exhibiting a faster regenerative capacity in comparison to larger animals (15-20 cm)^50^.

#### 4.2.2. Sample preparation

The specimens were initially fixed overnight at 4°C in a 4% paraformaldehyde solution. Following fixation, a triple wash with 1X PBS for 5 minutes each was conducted to remove residual fixative. Subsequently, the samples were immersed in a 30% sucrose solution in deionized water until they sank. To ensure optimal preservation of tissue structure, the samples were then embedded in Optimal Cutting Temperature (OCT) solution and stored at −80°C, maintaining their integrity for subsequent sectioning. The samples were cryo-sectioned at thickness between 20-30 µm and mounted on supercharge plus microscope slides. They were then baked at 65^°^C and stored at −80°C until further use.

#### 4.2.3. HCR-FISH probe design

The design of probes to detect *Trpv4*, *Piezo1*, and *Piezo2* mRNA followed the outlined procedures by Lovely et al. (2023) ^51^. In brief, the probe pairs were strategically designed with a two-base separation and included half of an HCR initiator sequence at the 3’ end. To facilitate this process, a web application called Probegenerator^51^ was used (http://ec2-44-211-232-78.compute-1.amazonaws.com/). It incorporates alignment to the axolotl genome for the creation of specific probe pools. The app leverages Oligominer^52^ and Bowtie2^53^ to generate probe set pools. The specific target sequences are used for each gene are listed in supplementary material section 7.

For fluorescent labeling, hairpins were sourced from Molecular Instruments (Los Angeles, CA). Specifically, *Trpv4*, paired with a B1 initiator, was matched with 600 pmol of B1 Alexa Fluor 647. Similarly, *Piezo1*, featuring a B2 initiator, was coupled with 600 pmol of B2 Alexa Fluor 488, and *Piezo2*, with a B3 initiator, was associated with 600 pmol of B3 Alexa Fluor 488/647 (*Trpv4* and *Piezo2* were analyzed on one section while *Piezo2* and *Piezo1* were analyzed on a different section). This tailored approach ensures precise pairing of initiators with the respective fluorophores for accurate detection without significant overlap in the excitation and emission spectrum^51^.

#### 4.2.4. mRNA Staining and Imaging

Following the procedures in Lovely et al. (2023) ^51^, the sections were washed with 1X PBS for 2 cycles of 5 minutes each. Subsequently, the sections were pre-hybridized with pre-warmed hybridization buffer (Molecular Instruments, Los Angeles, CA) at 37°C for 15 minutes, followed by an overnight incubation with probes (1:200 dilution in pre-warmed hybridization buffer) at 37°C in a humidified chamber.

On the subsequent day, the sections were subjected to 3 cycles of a 15-minute wash with pre-warmed formamide probe wash (Molecular Instruments, Los Angeles, C) at 37°C and a 10-minute wash (twice) with 5X saline sodium citrate buffer with 0.1% Tween (SSCT) at 37°C. H1 and H2 hairpins were aliquoted into distinct tubes (3 µM; Molecular Instruments, Los Angeles, C). They were subjected to 95°C temperature in a thermal cycler (Bio-Rad, Hercules, CA) for a minute and a half. Subsequently they were allowed to cool for a minimum of 30 minutes at room temperature in darkness.

The cooled H1 and H2 hairpin solutions were combined (2 µl in total) and prepared to a concentration of 1:50 by adding 48 µl of amplification buffer (Molecular Instruments, Los Angeles, C). For multiplexed studies with different hairpins, the amplification buffer volume was adjusted to maintain the dilution ratio at 1:50. After washing the sections with amplification buffer at room temperature for 10 minutes, they were incubated in the prepared hairpin solution with parafilm covering, overnight at room temperature. Subsequently, the sections were washed with 5X SSCT (3 times; 15 minutes each) at room temperature. The sections were then stained with DAPI nuclear stain (1:5000), washed with PBS, followed by DEPC water, and finally, mounted in Slow-fade™ with 1.5 microscope cover glass. To maintain RNA integrity within the tissue throughout the procedure, we consistently used RNAse-away spray^51,54^.

The samples were imaged using a confocal microscope (LSM 880, Zeiss, Germany) under the 20X objective and using AIRYSCAN fast mode. For each sample, 10 optical sections were acquired within the tissue section. After image acquisition, initial processing was performed in Zen Black software using the automatic 2D Airyscan setting. A maximum intensity projection was then generated^55^.

Background fluorescence of the images in each channel was subtracted using a median filter of size 20 pixels on ImageJ and overlaid on the nuclear stain. The mRNA expression was compared against negative controls which omitted the primary probes and contained only secondary fluorescent hairpins. To enhance visualization, each of the three image channels was processed using an inverted grayscale lookup table, and the images were cropped to highlight a smaller region of interest.

### 4.3. Immunofluorescence

The prepared slides, following the procedure outlined in 4.2.2, underwent blocking with 1.5% goat serum for 1 hour at room temperature. Subsequently, they were incubated overnight at 4°C with a mixture of 1:200 Rabbit Anti-TRPV4 polyclonal antibodies (antibodies-online Inc, Pottstown, PA) and 1:300 Rabbit Anti-PIEZO2 polyclonal antibodies (antibodies-online Inc, Pottstown, PA), prepared in 1.5% goat serum. Negative controls were treated with 1.5% goat serum instead of primary antibodies. Similar to HCR-FISH, mature chondrocytes in articular cartilage served as a positive control.

Following antibody incubation, the samples underwent three 5-minute washes with 1X PBS. Subsequent staining involved using 1:400 Goat Anti-Rabbit Alexa Fluor® 647 secondary antibodies, with negative controls subjected to the same secondary antibodies for 1 hour at room temperature. After three additional PBS washes for 5 minutes each at room temperature, the slides were stained with 1:5000 DAPI nuclear stain for 5 minutes at room temperature. Post-staining, the samples underwent two PBS washes and one deionized water wash for five minutes each at room temperature before being mounted in Slow-fade™ with 1.5 microscope cover glass. The imaging of the samples was conducted using a Zeiss LSM880 confocal microscope at 20X magnification.

The fluorescence intensity of mechanosensitive ion channels in blastemal tissues was standardized across all samples for a specific fluorophore threshold. Subsequently, a comparative analysis was performed between the fluorescence intensities across timepoints, considering both negative and positive controls for reference. All image processing was performed using ImageJ.

### 4.4. Calcium Signaling

Humerii from 9 fully matured axolotls (15-20 cm in length, n=3 humerii/group), including the cartilaginous distal epiphysis, were isolated and stained with a calcium indicator dye Calbryte® 520-AM (AAT-Bioquest, Pleasanton CA) in 1X Leibovitz’s L-15 medium at a concentration of 5µM at room temperature for 1 hour. After washing the samples with fresh 1X L-15 buffer solution, samples were then incubated for 30 minutes in either a DMSO vehicle control (0.05% v/v of DMSO), 10 µM GSK205 (MedChemExpress, Monmouth Junction, NJ), or 10 µM GdCl3 (Sigma Aldrich, St. Louis, MO) prepared in 1X L-15 media. They were then transferred to an imaging chamber containing the respective vehicle or antagonist solution. Calcium dynamics within the epiphysis were recorded for 10 minutes using a Zeiss LSM 880 confocal microscope equipped with a 20X objective.

Following baseline imaging in 1X L-15 medium (baseline), the medium was replaced with 60% L-15 media in deionized water supplemented with the same vehicle or drug concentration, and imaging was repeated for an additional 10 minutes to capture the chondrocyte response to hypotonic stress.

Image analysis was performed using previously published protocols^11,56^. Briefly, cell fluorescence intensities were tracked using Trackmate plugin in ImageJ. Peaks in fluorescence intensities were analyzed using Peak Finder function in MATLAB 2022 (Mathworks, Naitick, MA) and a cell was considered responsive if the peak prominence was at least four times that of baseline noise with a minimum peak width of 20s. 100-300 cells were tracked per sample.

Two-way repeated measures ANOVA followed by post-hoc analysis with Tukey correction (normal distribution and equal variances) was performed using GraphPad 10.3.0 (PRISM, San Diego, CA) to compare the percentage of cells exhibiting calcium signaling across treatment groups. Statistical significance was set at α < 0.05.

### 4.5. In-vivo Antagonist Treatments

#### 4.5.1. Animal Handling & Sample Collection

The forelimbs of 24 axolotls (∼3.5 cm in length) were amputated following the procedures in section 2.2. The animals were randomly assigned to either DMSO controlled vehicle, GSK205 or GdCl_3_ group. Treatment was administered by supplementing the housing water with the respective drug concentrations (GSK205 at 10 µM, GdCl3 at 10 µM or DMSO controlled vehicle) for a period of 2.5 weeks. The drug-infused water was changed 3 times a week to ensure sustained exposure to the treatments. During this period, the animals were monitored for general health, and the regenerating blastema were imaged using a brightfield microscope (Leica M165 FC) at 9, 20, and 27 DPA.

At the 27 DPA time point, the animals were sacrificed using 0.05% benzocaine followed by pithing^57^. Limbs from the animals were collected for alcian blue staining.

#### 4.5.2. Sample Processing and Analysis

The limbs collected from the animals at 27 DPA were fixed in 4% paraformaldehyde and stained with alcian blue following previously published protocols^58^. The stained limbs were imaged using a Zeiss Axio Zoom stereo microscope equipped with a 0.5X objective at a 5.4X zoom level (Zeiss, Germany). Total limb length and humeral length were quantified across treatment groups using ImageJ. Statistical analysis was performed using GraphPad 10.30.0 (PRISM, San Diego, CA), with a one-way ANOVA followed by a Dunnet’s post-hoc test (normal distribution, equal variances) to assess differences in total and humeral length among the treatment groups with respect to the control. An alpha value of less than 0.05 was considered significant.

## Acknowledgment

This project was funded by NSF grant # 2318594. Imaging was performed at the Center for Imaging Living Systems (CILS), Northeastern University. Portions of the manuscript are adapted from the author’s doctoral thesis (Kondiboyina, 2024)^44^.

## Author Contributions

Study design: VK, JM and SS; Data collection: VK,MM, MR, and AO; Data analysis and interpretation: VK, MM, MR, AO, TD, JM, and SS; Drafting of manuscript: VK, MM and SS; Revising manuscript: VK, MM, TD, JM and SS; Funding acquisition: SS, JM. SS takes responsibility for the integrity of the data analysis.

