## Supplementary Material for "TRPV4 Inhibition Reduces Cartilage Growth During Axolotl Limb Regeneration"

#### 1. Cumulative Gene Expression of Mechanosensitive Ion Channels

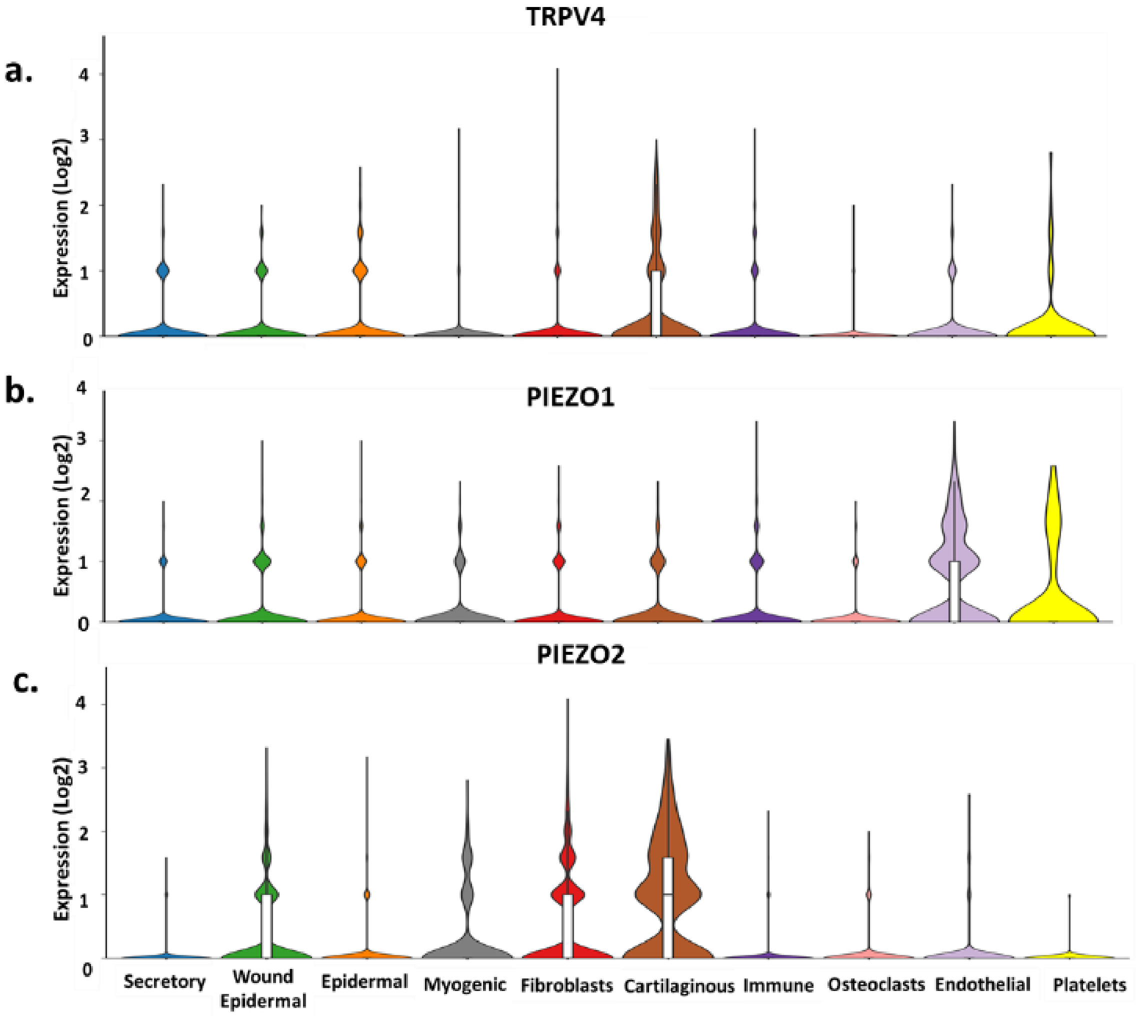


Figure S1. Cumulative gene expression in limb tissues across all regeneration timepoints (a) *Trpv4* (b) *Piezo1* (c) *Piezo2*. *Piezo1* has minimal expression in fibroblasts and cartilage cells. The central line or box plot inside the violin represents the median and interquartile range, while the width of the violin represents the density of the data at different values and the tails of the violin show the range of the data.

### 2. *Trpv4* and *Piezo2* HCR-FISH Negative Controls

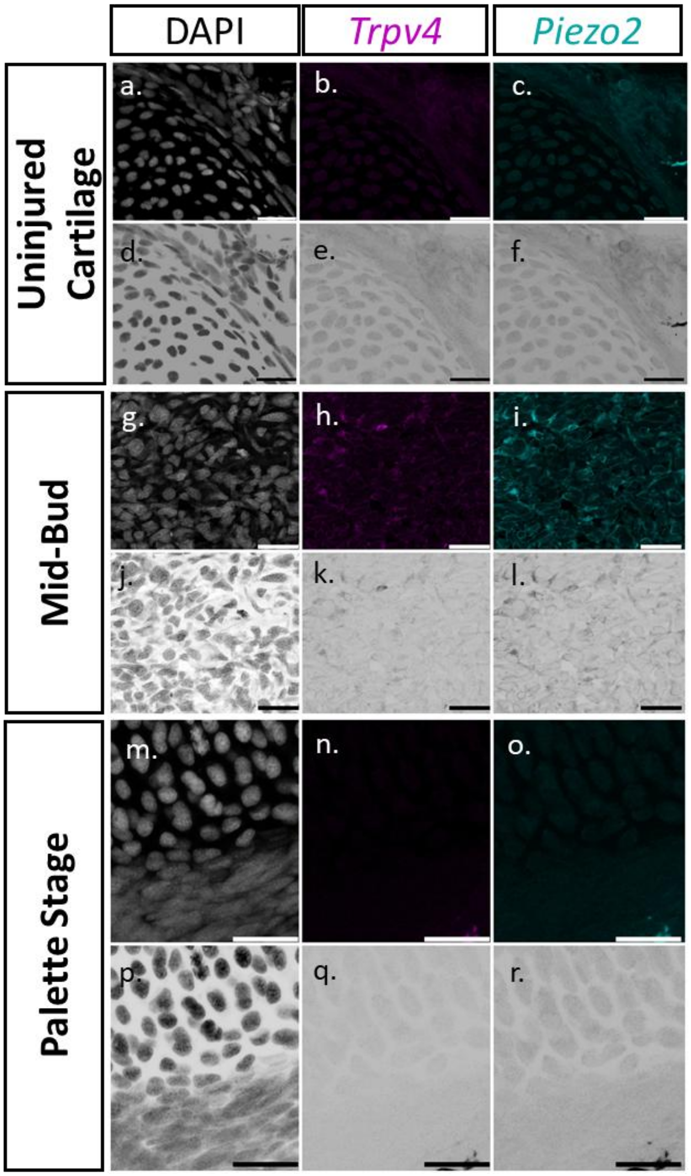


Figure S2. mRNA expression of *Trpv4* and *Piezo2* in negative controls (no primary probes, only secondary fluorescent hairpins) during limb regeneration at (a-f) Uninjured Cartilage (g-l) Mid-bud blastema (m-r) Palette stage blastema. Scale: 50µm. There was no discernable fluorescence signal in the *Trpv4 and Piezo2 channels*. The individual channels in the ROI are visualized as inverted grayscale lookup table images with the darker dots representing respective mRNA while the lighter circular boundary representing the cell nuclei.

#### 3. *Piezo1* and *Piezo2* HCR-FISH Negative Controls

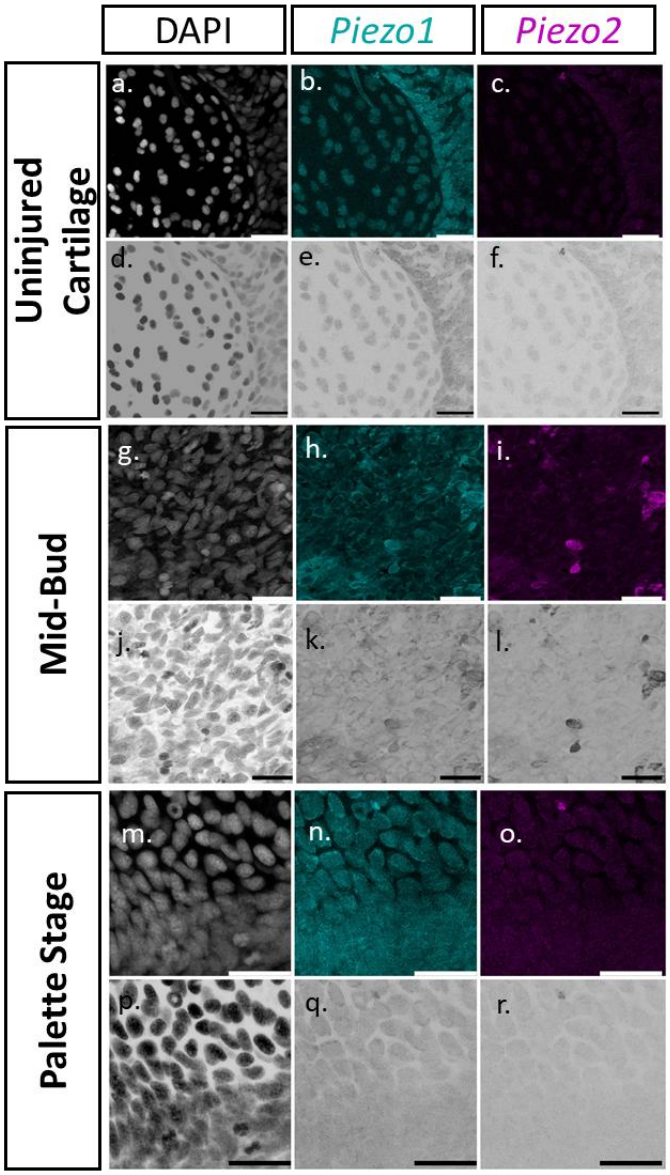


Figure S3. mRNA expression of *Piezo1* and *Piezo2* in negative controls (no primary probes, only secondary fluorescent hairpins) during limb regeneration at (a-f) Uninjured Cartilage (g-l) Mid-bud blastema (m-r) Palette stage blastema. Scale: 50μm. There was no discernable fluorescence signal in the *Piezo1 and Piezo2 channels*.

### 4. HCR-FISH *Piezo1and Piezo2 Expression*

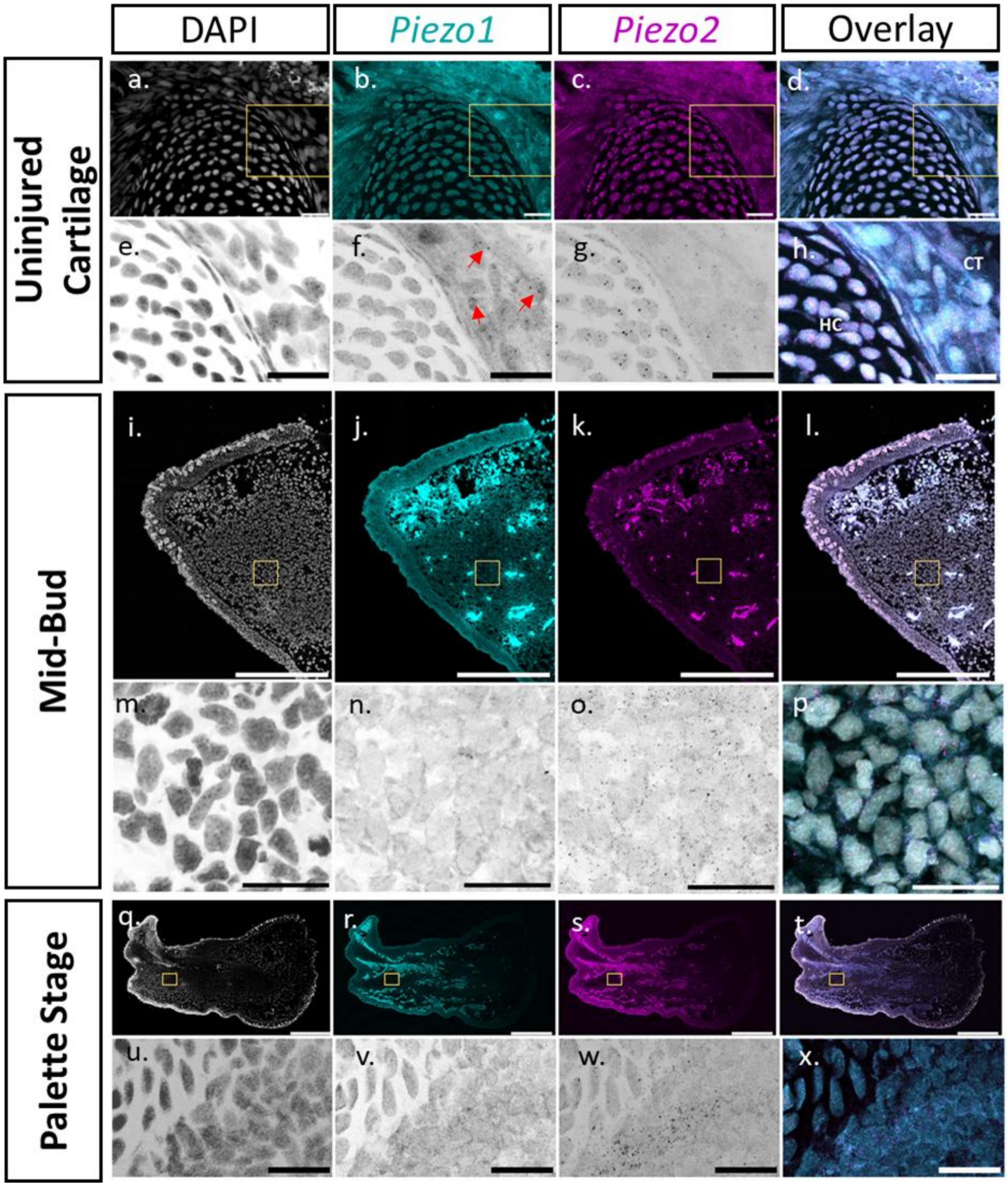


Figure S4. mRNA expression of *Piezo1* and *Piezo2* during limb regeneration at (a-h) Uninjured Cartilage. Scale (a-d) 500µm (e-h): 50µm (i-p) Mid-bud blastema. Scale (i-l) 1000µm, (m-p) 50µm and (q-x) Palette stage blastema. Scale (q-t) 1000µm (u-x) 50µm. CT: Connective Tissue. HC: Humeral Cartilage. Red arrows point to *Piezo1* expression.

### 5. HCR-FISH *Epithelium*

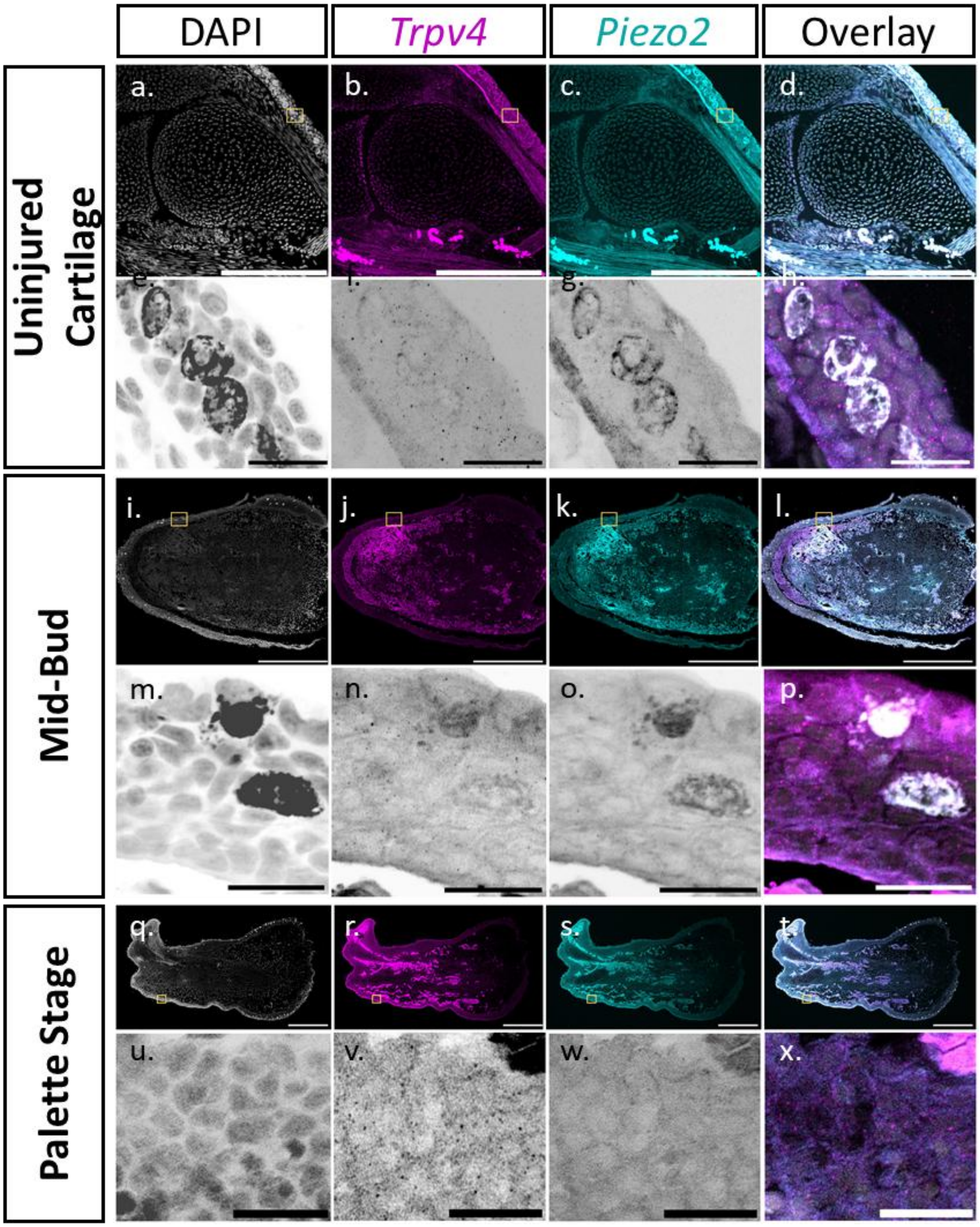


Figure S5. mRNA expression of *Trpv4* and *Piezo2* during limb regeneration in the epithelium (yellow ROI) at (a-h) Uninjured Cartilage (i-p) mid bud blastema stage (q-x) palette . *Trpv4* has robust expression in the epithelium at all regeneration timepoints. Scale: 50µm

### 6. Immunofluorescence Negative Controls

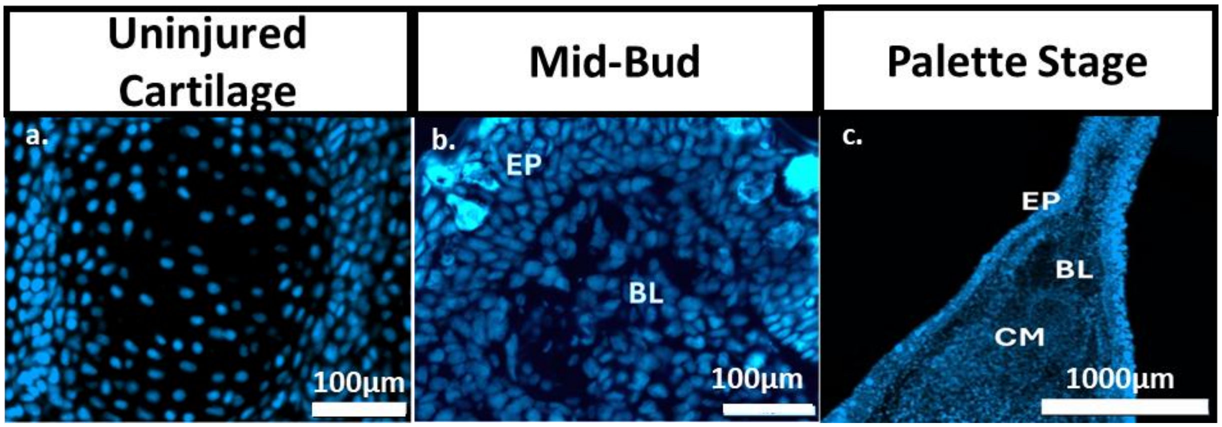


Figure S6. Negative controls for Goat Anti-Rabbit Alexa Fluor® 647 secondary antibodies used to evaluate TRPV4 and PIEZO2 protein expression during limb regeneration at (a) Uninjured cartilage (b) Mid-Bud Stage (c) Palette Stage. Cyan stains for the nucleus and magenta stains for the presence of secondary fluorescent antibodies. There was no discernable fluorescence signal in the antibody channel. EP: Epithelium, BL: De-differentiated Blastemal cells, CM: Condensed Mesenchyme

### 7. HCR-FISH Target Sequences for *trpv4, piezo1 and piezo2* :

| **Pool name** | **Sequence** |
|---|---|
| Trpv4_B1 | gAggAgggCAgCAAACggAACGCCAACATCAATAGAGCTGTTGAG |
| Trpv4_B1 | CGGCATTTCCATCATCAATCTGGGATAgAAgAgTCTTCCTTTACg |
| Trpv4_B1 | gAggAgggCAgCAAACggAAAAGAGGACAGCGGGAATGATTCGTT |
| Trpv4_B1 | CATCTTCATTCTCAAACAAATTGGCTAgAAgAgTCTTCCTTTACg |
| Trpv4_B1 | gAggAgggCAgCAAACggAATCCGTGTTGGCTCGATGGCCGGTGT |
| Trpv4_B1 | GCTTGTTGTCCCCGGCCGCGTGTGGTAgAAgAgTCTTCCTTTACg |
| Trpv4_B1 | gAggAgggCAgCAAACggAAATCCATTGGGTTAGGCATGCCTTTT |
| Trpv4_B1 | AGAATCGTAGATGGTGGATTCAAGATAgAAgAgTCTTCCTTTACg |
| Trpv4_B1 | gAggAgggCAgCAAACggAATGAATCCATGGGTGCTTTCCTGGGG |
| Trpv4_B1 | ATGGTGATAGGTCTCATAATCAAACTAgAAgAgTCTTCCTTTACg |
| Trpv4_B1 | gAggAgggCAgCAAACggAAGCGTCTTCGCTTGTTCTCTGTAGGG |
| Trpv4_B1 | ACTCTGTTTGTCTGCATTGATCTTTTAgAAgAgTCTTCCTTTACg |

| | |
|---|---|
| Trpv4_B1 | gAggAgggCAgCAAACggAAAGGCGGGTCTGGCGCTTGTGCTTTT |
| Trpv4_B1 | GTGCCGGTTGAACACCTTGATGACGTAgAAgAgTCTTCCTTTACg |
| Trpv4_B1 | gAggAgggCAgCAAACggAATCCGTCCAGGTCTGACGTCGACCCC |
| Trpv4_B1 | CTTCTGCGTCAACAGGTAAGTAAGCTAgAAgAgTCTTCCTTTACg |
| Trpv4_B1 | gAggAgggCAgCAAACggAATCGAAAATCCTCATCCGTCAGGCGT |
| Trpv4_B1 | TAAACATGTCTTCCCCGTCGACGCCTAgAAgAgTCTTCCTTTACg |
| Trpv4_B1 | gAggAgggCAgCAAACggAATCCGCTCAGGTTCATCAACGCTTTT |
| Trpv4_B1 | CAGCATCGGAATGGTGTCATTCTTCTAgAAgAgTCTTCCTTTACg |
| Trpv4_B1 | gAggAgggCAgCAAACggAATGAACTCTCGCAAGTTGCCTGTTTT |
| Trpv4_B1 | AGTAGACATCCCGAAACGGAGCATTTAgAAgAgTCTTCCTTTACg |
| Trpv4_B1 | gAggAgggCAgCAAACggAACAATGTGCAGCGCCGTCTGACCTCG |
| Trpv4_B1 | CATAATGCTTGCAGCGCCTCTCGACTAgAAgAgTCTTCCTTTACg |
| Trpv4_B1 | gAggAgggCAgCAAACggAACGGCTCCTTTCTCCACCAGCAGCTC |
| Trpv4_B1 | AGCGTCCGCGGGCCTGCGCGTGTACTAgAAgAgTCTTCCTTTACg |
| Trpv4_B1 | gAggAgggCAgCAAACggAAAGCCGCCCTCATCCCGGGGCTGGAA |
| Trpv4_B1 | ACAGGGGCAATTCACCAAAGTAGAATAgAAgAgTCTTCCTTTACg |
| Trpv4_B1 | gAggAgggCAgCAAACggAACCGGTTGGTTGGTGCAGGCAGCCAG |
| Trpv4_B1 | CATTTTCCGTGAGGTAGTGAACAATTAgAAgAgTCTTCCTTTACg |
| Trpv4_B1 | gAggAgggCAgCAAACggAATCCTGACGACGCAAGTCTGCCTTTT |
| Trpv4_B1 | GCATGGAGGACAGTGTTGCCGCGGGTAgAAgAgTCTTCCTTTACg |
| Trpv4_B1 | gAggAgggCAgCAAACggAAAAGAGATCATACATTTTCGTCAAAA |
| Trpv4_B1 | GGGAAAAGTTTGACACATTTGATCATAgAAgAgTCTTCCTTTACg |
| Trpv4_B1 | gAggAgggCAgCAAACggAATTGAGAAATGATTCCAGGCTGTCTC |
| Trpv4_B1 | ATCATCAGAGGGGAGAGTCCGTCATTAgAAgAgTCTTCCTTTACg |
| Trpv4_B1 | gAggAgggCAgCAAACggAAAGCCCAATTTTGCCCAACTTGGCCG |
| Trpv4_B1 | ATCTCCCTGCGTATGATATGCTGGATAgAAgAgTCTTCCTTTACg |
| Trpv4_B1 | gAggAgggCAgCAAACggAAGATAAATGGCGTGCATCCTCGTCCT |
| Trpv4_B1 | CCATAGGCCCAGTCTCTGAACTTCCTAgAAgAgTCTTCCTTTACg |
| Trpv4_B1 | gAggAgggCAgCAAACggAAAGGTCATAGAGAGAAGAGTAGACTG |
| Trpv4_B1 | TCCTCTCCACAGGTGTCCAATGATGTAgAAgAgTCTTCCTTTACg |
| Trpv4_B1 | gAggAgggCAgCAAACggAATACACCAAGATCTCCAGCACAGACA |
| Trpv4_B1 | TCATGGCGATTCTCTATCTTACTGTTAgAAgAgTCTTCCTTTACg |
| Trpv4_B1 | gAggAgggCAgCAAACggAATCATTGATCGGCTCCACCGCTAACA |
| Trpv4_B1 | AACTTCCGCCATTTGTCCCTCAGAATAgAAgAgTCTTCCTTTACg |
| Trpv4_B1 | gAggAgggCAgCAAACggAAACACTGATGTAGAAAGACACAGCTC |
| Trpv4_B1 | ATGATCATGGCGATGAGATAGGAAATAgAAgAgTCTTCCTTTACg |
| Trpv4_B1 | gAggAgggCAgCAAACggAAGGCCTGTAATATGCAGTCAATGTGA |
| Trpv4_B1 | TAAGGGTAAGGAGGGATGCCCTCCATAgAAgAgTCTTCCTTTACg |
| Trpv4_B1 | gAggAgggCAgCAAACggAAAGCCGCAGGTAATCTATCGTGGTGG |
| Trpv4_B1 | GTAAACAAAGTAATGATCTCTCCAGTAgAAgAgTCTTCCTTTACg |
| Trpv4_B1 | gAggAgggCAgCAAACggAAAAGAGGTCTTTGACATTTGTGAAAA |
| Trpv4_B1 | GAGTTCACCCCAGGACACTTCTTCATAgAAgAgTCTTCCTTTACg |

| | |
|---|---|
| Trpv4_B1 | gAggAgggCAgCAAACggAAAACTGAAATGAGCCGTCAATAAATA |
| Trpv4_B1 | ACCAGCACCGAATAAATGAAGTACATAgAAgAgTCTTCCTTTACg |
| Trpv4_B1 | gAggAgggCAgCAAACggAAGCCAGATACAGCGCTGCGGTGACGA |
| Trpv4_B1 | ATGACGGCGAGGTAAGCTTCAATGCTAgAAgAgTCTTCCTTTACg |
| Trpv4_B1 | gAggAgggCAgCAAACggAAATCCAACCAAGGACCAGGGCAAAGA |
| Trpv4_B1 | AACCCTCGAGTGAAGTAAAGTGTGTTAgAAgAgTCTTCCTTTACg |
| Trpv4_B1 | gAggAgggCAgCAAACggAAATGATGCTGTACGTTCCGGTCAGCT |
| Trpv4_B1 | AGGTCTTTAAAGAGGATCTTCTGGATAgAAgAgTCTTCCTTTACg |
| Trpv4_B1 | gAggAgggCAgCAAACggAAAGCAGGTACACCAACAGGAAGCGGA |
| Trpv4_B1 | AAAGCTGAGGCATAACCAATCATGATAgAAgAgTCTTCCTTTACg |
| Trpv4_B1 | gAggAgggCAgCAAACggAACTGGGACAAGGGTTCAGTAAAGACA |
| Trpv4_B1 | GTGGAGTTGTCGGTGCAGTTTTGTTTAgAAgAgTCTTCCTTTACg |
| Trpv4_B1 | gAggAgggCAgCAAACggAAGAGGGGTACTCAGGGGTTGGGCAGT |
| Trpv4_B1 | TTGCTGAAGGTGCTGCTGTCCCGACTAgAAgAgTCTTCCTTTACg |
| Trpv4_B1 | gAggAgggCAgCAAACggAAGTCAACTTGAAGAGGTCTAGAAGGA |
| Trpv4_B1 | ATCATCTCCAGATCGCCCATACCAATAgAAgAgTCTTCCTTTACg |
| Trpv4_B1 | gAggAgggCAgCAAACggAAAAGACTGCTGGGTATTTGGCACTGT |
| Trpv4_B1 | ATGATGTAGGTCACCAGCAGGATAATAgAAgAgTCTTCCTTTACg |
| Piezo1_B2 | CCTCgTAAATCCTCATCAAAACAGGCCCAAGCTGATGGCGGACAC |
| Piezo1_B2 | CAGATCTTTTCGGTATGAGGCGGTTAAATCATCCAgTAAACCgCC |
| Piezo1_B2 | CCTCgTAAATCCTCATCAAAGGATTCAATCGGTCTGCCTTTGGGG |
| Piezo1_B2 | TGCCTCTTCCTGCTCGAGTGGGTCCAAATCATCCAgTAAACCgCC |
| Piezo1_B2 | CCTCgTAAATCCTCATCAAACCTCCGATCTGGATCTGCAGGGTCC |
| Piezo1_B2 | GTCCTCCCCACCTGGGTGACCAAGGAAATCATCCAgTAAACCgCC |
| Piezo1_B2 | CCTCgTAAATCCTCATCAAATGCCCGATTAGGAGCCGCGGATTCA |
| Piezo1_B2 | CACACGCAGCCGAGCTTTCAGCAGCAAATCATCCAgTAAACCgCC |
| Piezo1_B2 | CCTCgTAAATCCTCATCAAACACTGTTTGGAGGACCTTGTGCGCG |
| Piezo1_B2 | AAGAAGAACGATGGCCAGAGACTTCAAATCATCCAgTAAACCgCC |
| Piezo1_B2 | CCTCgTAAATCCTCATCAAATGAGGGCATGATGATGCCTGCCAAC |
| Piezo1_B2 | GAGAAGGTAGTAAGCTCCAGAAAAGAAATCATCCAgTAAACCgCC |
| Piezo1_B2 | CCTCgTAAATCCTCATCAAACGCCCACCAGGTGCAGACTCCAATG |
| Piezo1_B2 | GCCCAGGTGGCTGATGGGAAAGTGGAAATCATCCAgTAAACCgCC |
| Piezo1_B2 | CCTCgTAAATCCTCATCAAAGCCCACCATCACGCACAACGCGTTG |
| Piezo1_B2 | ACAGATCAGGTGGCCACCAGCGAAGAAATCATCCAgTAAACCgCC |
| Piezo1_B2 | CCTCgTAAATCCTCATCAAAGAAGAACACGGCCTGGTAGCAGTAA |
| Piezo1_B2 | GAGGCTTCCAGATGGGATGAGCTGCAAATCATCCAgTAAACCgCC |
| Piezo1_B2 | CCTCgTAAATCCTCATCAAAATTCTTCAGGCCAAATAACCTGGCC |
| Piezo1_B2 | GCTGGTGCAGTTTGTGTACAGAATGAAATCATCCAgTAAACCgCC |
| Piezo1_B2 | CCTCgTAAATCCTCATCAAAGCTCACGTTGAGGACAAGGTCATTG |
| Piezo1_B2 | GGGGTTCACGTAGACTGGCCAATCAAAATCATCCAgTAAACCgCC |
| Piezo1_B2 | CCTCgTAAATCCTCATCAAAGGTGAAGTACAGAAGCAGCAGGACC |
| Piezo1_B2 | TTTCCTGAGCTTCAGCAGAGAGGCCAAATCATCCAgTAAACCgCC |

**TRPV4 Inhibition Reduces Cartilage Growth During Axolotl Limb Regeneration**

Vineel Kondiboyina, Melissa Miller, Maren Ritterbuck, Ashlin Owen, Timothy J Duerr, James R Monaghan, Sandra J Shefelbine

| | |
|---|---|
| Piezo1_B2 | CCTCgTAAATCCTCATCAAATAGCATTGGCTGGGCACTGGGCTCC |
| Piezo1_B2 | CGATCCTCCTGCGTCTAGAGCAGCAAAATCATCCAgTAAACCgCC |
| Piezo1_B2 | CCTCgTAAATCCTCATCAAACTCAGGTACAGCACTGACTTCTGCA |
| Piezo1_B2 | CTGATCTTGTTGGTGGTGGAATCCTAAATCATCCAgTAAACCgCC |
| Piezo1_B2 | CCTCgTAAATCCTCATCAAACTTCTGTGGCTCAGTCTTCTTCCGA |
| Piezo1_B2 | GCCTAACATGTGCAGAGGGGATTTCAAATCATCCAgTAAACCgCC |
| Piezo1_B2 | CCTCgTAAATCCTCATCAAAAATGTAGCTCTGGTTCATGATCACA |
| Piezo1_B2 | CCAGACCATCATGGCAATGAGGGCAAAATCATCCAgTAAACCgCC |
| Piezo1_B2 | CCTCgTAAATCCTCATCAAACGTCAGCCAGCTGTGGTAAGTGATG |
| Piezo1_B2 | GAGGCAGGCCCAAAGCAGCAACACGAAATCATCCAgTAAACCgCC |
| Piezo1_B2 | CCTCgTAAATCCTCATCAAACTGGCGCCGGTTCCGCACGATCCAG |
| Piezo1_B2 | AATGAAGGGCGAGCAGAGCATGGCAAAATCATCCAgTAAACCgCC |
| Piezo1_B2 | CCTCgTAAATCCTCATCAAACAGCTCGGGCTCGAGGTCCATCCCC |
| Piezo1_B2 | CAGGCTCATGAAGCCGATCCTCGTGAAATCATCCAgTAAACCgCC |
| Piezo1_B2 | CCTCgTAAATCCTCATCAAATTTTGCCCGAATAAGGCCCAGTTGC |
| Piezo1_B2 | CATGGCGCCCAGGTGGAAGCAGGGGAAATCATCCAgTAAACCgCC |
| Piezo1_B2 | CCTCgTAAATCCTCATCAAACAGCCAGAAGGTGAGAGTGTAGAGA |
| Piezo1_B2 | CAGCTCCTTCACAAACTGTCGCAACAAATCATCCAgTAAACCgCC |
| Piezo1_B2 | CCTCgTAAATCCTCATCAAAAGATGTCCTCCGCTTCCGTTTCAGG |
| Piezo1_B2 | TGCCACGGCCACCTCTGTGAGCGGCAAATCATCCAgTAAACCgCC |
| Piezo1_B2 | CCTCgTAAATCCTCATCAAAGTTCTGGTTTCTCTTGGGCCCTGCG |
| Piezo1_B2 | GACCATCGCTCCCAAGATCTTTAGAAAATCATCCAgTAAACCgCC |
| Piezo1_B2 | CCTCgTAAATCCTCATCAAACATCCAGTACTTGGCGTAGAAGCCC |
| Piezo1_B2 | CACGATGAACATTCCAGCGCAAACAAAATCATCCAgTAAACCgCC |
| Piezo1_B2 | CCTCgTAAATCCTCATCAAAGACAACCAGCCGCCCAGCAAAGCTG |
| Piezo1_B2 | GAATAGGAACATGTAAACTATCTTGAAATCATCCAgTAAACCgCC |
| Piezo1_B2 | CCTCgTAAATCCTCATCAAACACCTGGAAAAGTGTCAGGCACAGG |
| Piezo1_B2 | CAGCAGCTTCCGCCAGAGGGAGTAGAAATCATCCAgTAAACCgCC |
| Piezo1_B2 | CCTCgTAAATCCTCATCAAATAGCATGAGCACCAAGAGGTGATTC |
| Piezo1_B2 | GCGGTAAACAATTGCTTCAAAGACCAAATCATCCAgTAAACCgCC |
| Piezo1_B2 | CCTCgTAAATCCTCATCAAAATAACGCTTGCGGTAGTATTCCTGG |
| Piezo1_B2 | AGCCTGGGTGCTAGGTGGCAGAAGTAAATCATCCAgTAAACCgCC |
| Piezo1_B2 | CCTCgTAAATCCTCATCAAACTTCTCTCGTGTGACTTCTTCAAAG |
| Piezo1_B2 | GACACAGCTGACGACGCCCTGGTCTAAATCATCCAgTAAACCgCC |
| Piezo1_B2 | CCTCgTAAATCCTCATCAAAATCTCCAAGCCAAACTTGTAGAAAA |
| Piezo1_B2 | ATCACATTCACGGTCAGCAGGAAGCAAATCATCCAgTAAACCgCC |
| Piezo1_B2 | CCTCgTAAATCCTCATCAAAATAACCATGAAGTTCATGCGTTGGC |
| Piezo1_B2 | ATGACGATTAGCCAGCAGCCATGTAAAATCATCCAgTAAACCgCC |
| Piezo1_B2 | CCTCgTAAATCCTCATCAAAATGGCTCCTCGCCGGCGCCGTGTCA |
| Piezo1_B2 | AGGCAGTACTTTGGCCAGAGCTTGGAAATCATCCAgTAAACCgCC |
| Piezo1_B2 | CCTCgTAAATCCTCATCAAATGGTAGAGAATGAAGAAGAGGAGGA |
| Piezo1_B2 | GGCGGCATCCCCACACACAGCAGATAAATCATCCAgTAAACCgCC |
| Piezo1_B2 | CCTCgTAAATCCTCATCAAACTCCAAGGGTAGTCTATGCAGAGGA |

| | |
|---|---|
| Piezo1_B2 | GAGTTAATTGGTATTGAATGCTTCCAAATCATCCAgTAAACCgCC |
| Piezo1_B2 | CCTCgTAAATCCTCATCAAAGGCAGATACATCCACTTGATCAGAG |
| Piezo1_B2 | GTTGAATTTGGAGGAATGTAGAAATAAATCATCCAgTAAACCgCC |
| Piezo1_B2 | CCTCgTAAATCCTCATCAAAAGCAACAGGAAATCACTGATGAGGT |
| Piezo1_B2 | ACGTTCCACTGCTGGGAAGCACAAAAAATCATCCAgTAAACCgCC |
| Piezo2_B3 | gTCCCTgCCTCTATATCTTTTTGTCTTTGTCGGTTCCGAAAACAG |
| Piezo2_B3 | ATAACCGTCCTGTATGTCCGTGCATTTCCACTCAACTTTAACCCg |
| Piezo2_B3 | gTCCCTgCCTCTATATCTTTAAAAGCTGGAGAAACATATGGCTTT |
| Piezo2_B3 | GGAATATAATGTTAAGAAGTAGGAATTCCACTCAACTTTAACCCg |
| Piezo2_B3 | gTCCCTgCCTCTATATCTTTTGGCCTCGAGGCTGGCAATGGTGAT |
| Piezo2_B3 | AGTTAAAATTTGGTTCGATGTTGTTTTCCACTCAACTTTAACCCg |
| Piezo2_B3 | gTCCCTgCCTCTATATCTTTGCCTGAAGGACTTCTCCCACGTTGT |
| Piezo2_B3 | CACCTTTCATGCTTTCAAATCCTAATTCCACTCAACTTTAACCCg |
| Piezo2_B3 | gTCCCTgCCTCTATATCTTTACATTCGCAGTCCATTGCCAGCATC |
| Piezo2_B3 | CAATGAACATACCGATATCTGGCACTTCCACTCAACTTTAACCCg |
| Piezo2_B3 | gTCCCTgCCTCTATATCTTTTACAGACCAGCCACGTGGCCAGGCT |
| Piezo2_B3 | CTGTGAGTGTCTTCTGAGCGAGGTTTTCCACTCAACTTTAACCCg |
| Piezo2_B3 | gTCCCTgCCTCTATATCTTTATTGTGAGTTGTACTGTGCTGCATC |
| Piezo2_B3 | CTGCCACTGCCAGCTCCTCATTTTCTTCCACTCAACTTTAACCCg |
| Piezo2_B3 | gTCCCTgCCTCTATATCTTTTCATCACTTCATCCATCTCCATCTT |
| Piezo2_B3 | CTCCAGCATCCAGATCTTCTTCAAATTCCACTCAACTTTAACCCg |
| Piezo2_B3 | gTCCCTgCCTCTATATCTTTTCTCTTCATTCTCTTCTTCATCTTC |
| Piezo2_B3 | TCCTACGAAGTAGTTTTAGTTTTGTTTCCACTCAACTTTAACCCg |
| Piezo2_B3 | gTCCCTgCCTCTATATCTTTCTTTGAGTTTGGAAGCTATAAAAGC |
| Piezo2_B3 | CGGTGGTTATAATATTGCCAATTAGTTCCACTCAACTTTAACCCg |
| Piezo2_B3 | gTCCCTgCCTCTATATCTTTGTAAAATTGTCACTACAACTTTTCC |
| Piezo2_B3 | ATGGCAGCATCATACCTGTTACACCTTCCACTCAACTTTAACCCg |
| Piezo2_B3 | gTCCCTgCCTCTATATCTTTGACCACCATGTGCAGAGCCCCAAAA |
| Piezo2_B3 | ATCAGCGGATCAAATACTCGGCAGCTTCCACTCAACTTTAACCCg |
| Piezo2_B3 | gTCCCTgCCTCTATATCTTTGCCATTAATACACATAGACAGCTGA |
| Piezo2_B3 | CCAATCAAGTGCCCTGCACTGAATATTCCACTCAACTTTAACCCg |
| Piezo2_B3 | gTCCCTgCCTCTATATCTTTCTCCTCCTCTTCCTCTTCTTTCTTT |
| Piezo2_B3 | CTCTGTCTCATCTTGCTCTTCTTCCTTCCACTCAACTTTAACCCg |
| Piezo2_B3 | gTCCCTgCCTCTATATCTTTATGCACTTTAGAGTTGTCCCTTTCA |
| Piezo2_B3 | GATAAACTGGAAGACGGTGACCATTTTCCACTCAACTTTAACCCg |
| Piezo2_B3 | gTCCCTgCCTCTATATCTTTAAGGGCACAGATGTAGCTTTGCTTC |
| Piezo2_B3 | CGTGATGCTCCAGGCCATCATGGAATTCCACTCAACTTTAACCCg |
| Piezo2_B3 | gTCCCTgCCTCTATATCTTTGAGCACAAAGGTGAGCCAGCTGTGA |
| Piezo2_B3 | CATCCAAAGGGCGCAGGACCAGATGTTCCACTCAACTTTAACCCg |
| Piezo2_B3 | gTCCCTgCCTCTATATCTTTCATGGCGTACTTCCGCCTGTCGCGG |
| Piezo2_B3 | ATAGACGACCATGAACGGAGAACTATTCCACTCAACTTTAACCCg |
| Piezo2_B3 | gTCCCTgCCTCTATATCTTTGTACTGCAGCGTCACCAGCAGGTTT |

**TRPV4 Inhibition Reduces Cartilage Growth During Axolotl Limb Regeneration**

Vineel Kondiboyina, Melissa Miller, Maren Ritterbuck, Ashlin Owen, Timothy J Duerr, James R Monaghan, Sandra J Shefelbine

| | |
|---|---|
| Piezo2_B3 | GCTGTCGTTGAGGTCGATGCTCCAGTTCCACTCAACTTTAACCCg |
| Piezo2_B3 | gTCCCTgCCTCTATATCTTTTAGAAATCCACTGTACGCGCGCAGC |
| Piezo2_B3 | AGCAAGTTCTCCAGGAACCTTTCTTTTCCACTCAACTTTAACCCg |
| Piezo2_B3 | gTCCCTgCCTCTATATCTTTGAAAGTGATATTGAAAACAATCTTG |
| Piezo2_B3 | GGTGAGGTGTTGGCGGAGCAAAAGCTTCCACTCAACTTTAACCCg |
| Piezo2_B3 | gTCCCTgCCTCTATATCTTTTTCTTTCACCTTCTGCGCTTTCTGC |
| Piezo2_B3 | TCCTACTTTGACCTCGGATAATGTTTTCCACTCAACTTTAACCCg |
| Piezo2_B3 | gTCCCTgCCTCTATATCTTTGCTAGACTCCCATCTTGGTTTACCA |
| Piezo2_B3 | GCAGTTAGGTTCATCATGGTGATGTTTCCACTCAACTTTAACCCg |
| Piezo2_B3 | gTCCCTgCCTCTATATCTTTTGACTTCTCCTCAAGTCTCGTCCCC |
| Piezo2_B3 | ATCCTCCTCCTCCTCGTTCTGGTCATTCCACTCAACTTTAACCCg |
| Piezo2_B3 | gTCCCTgCCTCTATATCTTTCTCGTCTTCTCCATCCCCTCGGTCA |
| Piezo2_B3 | TTCATCGTCCTTTTCCTCTTTTCCTTTCCACTCAACTTTAACCCg |
| Piezo2_B3 | gTCCCTgCCTCTATATCTTTCTCCGATTCATCCTCGACCTCTTCC |
| Piezo2_B3 | CTTTAGATCTGAAGACTCCTCTTCTTTCCACTCAACTTTAACCCg |
| Piezo2_B3 | gTCCCTgCCTCTATATCTTTGCGGTCAATAACCAAATGCCACTTG |
| Piezo2_B3 | CAGGAATTTCAAGAACAGCACAGTATTCCACTCAACTTTAACCCg |
| Piezo2_B3 | gTCCCTgCCTCTATATCTTTCAGCTCCAAAATCCACCATACAAAA |
| Piezo2_B3 | GTACGAAGACACAATTTTGATAATGTTCCACTCAACTTTAACCCg |
| Piezo2_B3 | gTCCCTgCCTCTATATCTTTGACCTCTCTCACTGTGACCCAGATG |
| Piezo2_B3 | AATTACGAAGACATAGTTAAACAGATTCCACTCAACTTTAACCCg |
| Piezo2_B3 | gTCCCTgCCTCTATATCTTTGGAATAAGGCAGAGCAAAAGCCCAG |
| Piezo2_B3 | CACGCTTGATGCCAGTGGGCGGAACTTCCACTCAACTTTAACCCg |
| Piezo2_B3 | gTCCCTgCCTCTATATCTTTAATGATGACGCACGTCCAAACAGTG |
| Piezo2_B3 | TGTGAGTTGGTACAACATTTTACAGTTCCACTCAACTTTAACCCg |
| Piezo2_B3 | gTCCCTgCCTCTATATCTTTACTGGAGAAGCTGGGAGGATCAATA |
| Piezo2_B3 | TTGATTTTCCGAAGGCACTGTGCAGTTCCACTCAACTTTAACCCg |
| Piezo2_B3 | gTCCCTgCCTCTATATCTTTTAGCTCTGTTCTGTTTGCCACGTTG |
| Piezo2_B3 | GGGAGCACTGTAGAGCATGGACGTCTTCCACTCAACTTTAACCCg |
| Piezo2_B3 | gTCCCTgCCTCTATATCTTTCAATCCAACCCAGCCTGAAGGGTCA |
| Piezo2_B3 | GTATGCAAGCAGCGGCGATGACTTCTTCCACTCAACTTTAACCCg |
| Piezo2_B3 | gTCCCTgCCTCTATATCTTTGGCTAGCATCAGCAGATTATTCCTT |
| Piezo2_B3 | GTAGATTGTCACTTCGAAGGCGAGGTTCCACTCAACTTTAACCCg |
| Piezo2_B3 | gTCCCTgCCTCTATATCTTTTCGGCATCGGTAGTACTCTTGGTGC |
| Piezo2_B3 | TTTAGTCACAGGTGTGGTGAGGTTGTTCCACTCAACTTTAACCCg |
